# Moss BRCA2 lacking canonical DNA binding domain promotes efficient homologous recombination and binds to DNA

**DOI:** 10.1101/2025.03.19.644124

**Authors:** Alice Chanteau, Suliane Quilleré, Arthur Crouset, Sreejith Allipra, Ulysse Tuquoi, Pierre-François Perroud, Simona Miron, Pauline Dupaigne, Sophie Zinn-Justin, Fabien Nogué, Rajeev Kumar

## Abstract

BRCA2 interacts with RAD51 and DMC1 recombinases and binds to single-stranded DNA (ssDNA), through its canonical DNA binding domain (DBD) to mediate homology-directed DNA repair (HDR). While the well-folded DBD is widely conserved in diverse eukaryotes, a non-canonical BRCA2 variant lacking this domain is found in *Drosophila melanogaster*. Whether such a non-canonical BRCA2 variant exists in other species is unknown. Additionally, the DNA-binding activity of a BRCA2 variant lacking DBD remains unclear. Here, we identify a new non-canonical BRCA2 in the model plant *Physcomitrium patens* (PpBRCA2). We establish that PpBRCA2 is essential for genome integrity maintenance, somatic DNA repair, HDR-mediated gene targeting, and RAD51 foci recruitment at DNA break sites. PpBRCA2 is also critical for DNA repair during meiosis, but interacts only weakly with DMC1, suggesting a distinct meiotic function compared to other BRCA2 homologs. Despite lacking the canonical DBD, PpBRCA2 binds ssDNA through its disordered N-terminal region and efficiently promotes HDR. Our work highlights that the ssDNA binding capacity of BRCA2 homologs is conserved regardless of the presence of canonical DBD and provides a deeper understanding of BRCA2’s functional diversity across species.

**Three Key messages:**

- *Physcomitrium patens BRCA2* (PpBRCA2) lacking canonical DBD mediates homologous recombination and promotes genome stability
- PpBRCA2 differentially interact with RAD51 and DMC1
- PpBRCA2 binds single-stranded DNA via its N-terminal to compensate for the loss of canonical DBD

## INTRODUCTION

The BREAST CANCER gene 2 (BRCA2) preserves genomic integrity in most eukaryotes. In humans, mutations in the *BRCA2 gene* predispose to a high risk of developing breast, ovarian, and other types of cancers (1, 2). The loss of BRCA2 functions in various eukaryotic species exhibits a spectrum of genome instability traits including hypersensitivity to genotoxic agents, chromosomal breakage, and aberrations (3–12). BRCA2 plays diverse roles in genome maintenance, including protecting stalled replication forks, suppressing single-stranded DNA (ssDNA) gaps, repairing interstrand DNA crosslinks, and cytotoxic DNA double-strand breaks (DSBs) (1, 2, 13–17). BRCA2 is essential for the repair of accidental breaks or programmed DSBs during meiosis through homologous recombination, also called homology-directed repair (HDR). Alternatively, other pathways including Non-Homologous End Joining (NHEJ) can repair DSB (18).

BRCA2 is a crucial mediator of HDR in somatic and meiotic cells. During HDR, Replication Protein A (RPA) binds to 3’ ssDNA overhangs generated by nucleolytic processing of DSBs. Two eukaryotic structurally and functionally related recombinases - RAD51 and its meiosis-specific paralog DMC1-replace RPA to form nucleoprotein filaments (19, 20). Only RAD51 acts in somatic cells, while both RAD51 and DMC1 are necessary during meiosis. These nucleofilaments drive homology searches and strand invasion into an intact donor template, producing recombination intermediates that are subsequently resolved as crossover (CO) or non-crossover repair products (19). However, under physiological conditions, RAD51 and DMC1 cannot outcompete RPA for ssDNA due to RPA’s higher binding affinity (21, 22) and require various mediator proteins that facilitate the formation and activation of RAD51 and DMC1 nucleofilaments (23). BRCA2 nucleates and stabilizes recombinase nucleofilaments through its direct interaction with recombinases and binding to DNA (5, 24–28). With its partner DSS1, BRCA2 facilitates RPA displacement from ssDNA, enabling nucleoprotein filament assembly (22, 29). DSS1 attenuates RPA’s affinity for ssDNA through DNA mimicry (22). Consequently, BRCA2 and DSS1 are essential for forming RAD51 and DMC1 nuclear foci during HDR (4, 9, 30–32). Thus, BRCA2 interactions with both RAD51/DMC1 and DNA appear to be critical for its function.

BRCA2 interacts with RAD51 and DMC1 through highly conserved BRC repeats (∼35 amino acids) found in one or multiple copies across BRCA2 homologs (33). Human BRCA2 contains eight BRC repeats, each with a varying binding affinity to RAD51 and DMC1 (34–37). Each human BRC repeat has two conserved key tetrameric hydrophobic motifs: 1) the FxxA motif, which mimics the RAD51 oligomerization interface of RAD51 and interacts with its catalytic domain (38, 39). 2) an φφx[E/D] motif (φ = hydrophobic residue) in the α-helical region, which engages a different hydrophobic pocket of the RAD51’s catalytic domain (39). These two distinct motifs have opposing effects – stimulating and inhibiting nucleoprotein filaments *in vitro* –providing insights into how BRCA2 binds recombinase monomers or oligomers (39). Additionally, RAD51 and DMC1 can interact with BRCA2 motifs outside BRC repeats (24, 27, 40).

Most BRCA2 homologs contain a well-folded canonical ssDNA-binding domain (DBD) and additional auxiliary regions for ssDNA and/or dsDNA interaction (33, 41–43). They together collectively facilitate nucleofilament formation through direct binding to ssDNA or via a diffusion-assisted delivery mechanism (44). The canonical DBD is a well-structured and highly conserved region composed of a helical domain, three oligonucleotide-binding (OB) folds, and a tower domain (45). It also harbors many cancer-associated pathogenic mutations in humans (Julien et al. Biomolecules 2021). The small acidic protein DSS1 binds to canonical DBD in humans (29, 46) and restrains the dsDNA binding activity of DBD to ensure BRCA2 targeting to ssDNA (47). Surprisingly, mammalian cells with BRCA2 lacking the canonical DBD remain viable and capable of HDR, suggesting additional DBDs in mammalian BRCA2 may compensate for the loss of canonical DBD (48–51). Notably, the *Drosophila melanogaster* BRCA2 homolog (DmBRCA2) lacks the canonical DBD (3, 5), but it remains unclear whether DmBRCA2 binds to DNA at all.

While a flowering model plant *Arabidopsis thaliana* has two BRCA2 homologs (AtBRCA2A and AtBRCA2B) containing BRC repeats and a canonical DBD (9), no BRCA2 homolog was identified in moss genomes so far (52). *Physcomitrium patens* is a moss model plant from the bryophyte clade and is often called a “green yeast” due to its high gene-targeting (GT) efficiency, comparable to *Saccharomyces cerevisiae*, allowing precise genetic modifications via homologous recombination (53). Over the past few decades, *P. patens* has emerged as a powerful model for studying HDR, DNA repair, and genome editing (54). Here, we report the identification and functional characterization of the *P. patens* BRCA2 homolog (PpBRCA2). The PpBRCA2 contains BRC repeats but lacks the canonical DBD, as previously observed for *Drosophila melanogaster* BRCA2. We have delineated the mechanism of non-canonical PpBRCA2 in HDR, DNA repair, and genome stability, demonstrating its mediator roles and the DNA-binding activity despite the absence of a canonical DBD. Based on our findings, we propose that two classes of BRCA2 homologs exist in eukaryotes: canonical BRCA2 with DBD and non-canonical BRCA2 lacking DBD.

## MATERIAL AND METHODS

### Plant material and culture

The *Physcomitrium patens* accessions “Gransden” and “Reute” were used as wild-type references (Haas et al., 2020). Four *BRCA2* full deletion lines were generated using the CRISPR-Cas9 system with two guide RNAs targeting the start codon and 3′ untranslated region (UTR) of the Pp6C10_10830/*PpBRCA2* gene: *brca2Δ-1* and *brca2Δ-2* in the Reute ecotype, and *brca2Δ-3* and *brca2Δ-4* in the Gransden ecotype. The *rad51-1-2* double mutant was generated by targeting *PpRAD51-1* (*Pp6c11_11820*) and *PpRAD51-2* (*Pp6c7_5100*) genes by CRISPR-Cas9 system in the Reute background only. All guides and genotyping primers are listed in the supplementary DATASET_S3.

Moss strains were maintained and propagated on rich agar PpNH4 medium overlaid with cellophane or as spot inocula on minimal BCD medium for growth and sporogenesis assays (55, 56). Culture chamber conditions were set at 60% humidity with 16/8 h light/dark long-day photoperiod at 23 °C. Protoplast isolation, PEG-mediated transfection, and selection of transformants followed established protocols (57). The short-day photoperiod of 8/16h light/dark at 15°C was used for sporogenesis induction on the BCD medium (55). Young sporophytes fixed in ethanol-acetic acid (3:1) were analyzed for meiosis (58) and mature sporophytes were assessed for spore viability by staining with Alexander solution (59) and for spore germination rates (60). Statistical analyses and graphs were generated using Prism GraphPad 10.3.0.

### Genes identification

For moss BRCA2 identification, HHpred searches with full-length *A. thaliana* BRCA2A (At4g00020) were performed as described previously (61, 62). The *Pp6C10_10830* gene encoding PpBRCA2 without intron was cloned using *P. patens* genomic DNA with PCR primers listed in DATASET_S3. The PSI-BLAST searches using *A. thaliana* BRCA2A as a query and the Phytozome.v13 database identified BRCA2 orthologs from the following species: *Ceratopteris richardii* (Ceric.07G096700), *Marchantia polymorpha* (Mapoly0045s0096), *Micromonas pusilla* (MicpuC2.est_orfs.10_1849_4269375:2) and *Chlamydomonas reinhardtii* (Cre13.g566900_4532). *Anthoceros agrestis* BRCA2 (AagrOXF_EVM_utg000116l.287) was identified using a specific Anthoceros BLAST database (63) (https://www.hornworts.uzh.ch/en/Blast.html). Protein sequence alignments including BRC repeats of HsBRCA2 (HGNC:1101) and DmBRCA2 (FB: FBgn0050169) were performed using CLUSTAL Omega (64).

### Vector design, cloning, and gene expression

All PCR primers and sgRNAs are listed in DATASET_S3. The guide RNAs targeting *PpBRCA2* (*Pp6c10_10830*), *PpRAD51-1* (*Pp6c11_11820*), or *PpRAD51-2* (*Pp6c7_5100*) genes were designed using the CRISPOR 4.97 tool (https://crispor.gi.ucsc.edu/) and were cloned into sgRNA expression cassettes either in the pDONR207 background by Gateway^TM^ (Thermo Fisher Scientific) or in the pDGB3 background by GoldenBraid cloning (65, 66). For GFP fusion at the PpBRCA2 N-terminal, a donor matrix harboring the *eGFP* encoding sequence flanked by *PpBRCA2* 5’ UTR and coding DNA sequence (CDS) was synthesized by Twist Bioscience (San Francisco, CA, USA) and cloned into a pDONR207-NeoR vector by Gateway^TM^ BP reaction (Thermo Fisher Scientific). *PpBRCA2* guide RNA targeting the start codon was used for GFP insertion at the endogenous locus. The complementation studies were performed with the synthesized CDS of *PpBRCA2\** with polymorphisms*, CpBRCA2, AtBRCA2*, and *AtBRCA2^ΔDBD^* without DBD flanked with 5’ and 3’ UTR at the *PpBRCA2* locus and cloned into a pTwistAmpHighCopy plasmid by Twist Bioscience (San Francisco, CA, USA). The CDS encoding *PpBRCA2* was cloned in the pUbi-Gateway vector for transient expression in *P. patens* (60, 67). Through CRISPR-Cas9 and a sgRNA targeting the deleted *PpBRCA2* locus, DSB was generated to facilitate the insertion of *PpBRCA2*, CpBRCA2,* or *AtBRCA2^ΔDBD^* from the above-cited repair matrices, while transiently expressing *PpBRCA2*. Plants with insertions were allowed to grow without selection and the absence of pUbi-Nos-PpBRCA2 integration was monitored by PCR. For yeast-two hybrid assays, DNA sequences encoding *PpBRCA2*, *PpRAD51-1*, *PpRAD51-2*, *PpDMC1*, or *BRC* repeats were cloned in pGAD-T7 and pGBK-T7 vectors. The fidelity of all constructs was confirmed by sequencing. RT-PCR determined transcript abundance from cDNA prepared from respective genotypes as described in (58) with primers listed in DATASET_S3.

### Genotoxic sensitivity and mutator phenotype assays

UV-B sensitivity assays were performed according to (58). For the bleomycin sensitivity assay, approximately 2500 protoplasts per well in a 24-well plate were chronically exposed to indicated doses of bleomycin on an agar regeneration medium (Fig. 4B). The protoplast regeneration rate was quantified after 72h from three replicates in each dose. The frequency of spontaneous mutations at the *APT* locus in wild-type and *brca2Δ* mutants was analyzed as previously described (58).

### Gene targeting assay

The gene targeting efficiency at the *APT* locus in the wild-type and *brca2Δ* mutants was measured as described previously (Collonnier et al. 2017a). *P. patens* protoplasts were transformed with the following plasmids: pUbi-Cas9 (expressing Cas9) (68), psgRNA-PpAPT#2 (Expressing guide RNA to target *APT* locus), and PpAPT-KO7 (Donor matrix containing a G418 resistance cassette flanked by genomic *APT* sequences). The double resistance to 2-FA and G418 can be conferred by targeted integration of the PpAPT-KO7 cassettes at the *APT* locus or cassette integration elsewhere in the genome coupled with non-sense mutations at the *APT* locus. To estimate the HDR frequency versus non-HDR, transformants with the disrupted *APT* gene due to cassette insertion at the locus by HR or non-sense mutation via non-HDR pathways were first selected on PpNH4 supplemented with 10 μM 2-FA, followed by another round of selection on media with 50 mg l^−1^ G418 (Duchefa). The *APT* locus-specific insertion was confirmed by PCR analysis with primers amplifying insertion borders or single-copy insertion (69). The HDR frequency was calculated by estimating G418-resistant plants among 2-FA-resistant transformants. Independently from HDR frequency measurements, the ratio of targeted integrations versus random integrations was calculated from the number of stable G418 resistant transformants obtained after three successive rounds of selection on 50 mg/l G418 (Duchefa) followed by selection on 2-FA to identify *APT* locus-specific insertion. G418-resistant plants showing amplification of cassette but not at the *APT* locus by PCR were considered random integrations (Fig. S4 A) (69).

### Cytological techniques

Meiotic chromosome spreads were prepared as previously described (58). For RAD51 immunostaining, we adopted a method described for Arabidopsis root tip nuclei according to (70). Briefly, 6-day-old *P. patens* protonema filaments from WT or *brca2Δ* were incubated with or without 5 µg/ml bleomycin for 3h for DSB induction. *P. patens* filaments were first fixed for 45 min in 4% paraformaldehyde in PME [50 mM piperazine-N, N’-bis(2ethanesulphonic acid) (PIPES), pH 6.9; 5 mM MgSO4; 1 mM EGTA] followed by three washes in PME for 5 min. Then, filaments were digested for 30 min at 37°C in digestion buffer (1% (w/v) cellulase, 0.5% (w/v) cytohelicase, 1% (w/v) pectolyase in PME) and were washed three times for 5 min in PME. Digested cells were then squashed gently onto slides with a coverslip and immersed in liquid nitrogen. Slides were air-dried and stored at −80°C. RAD51 immunostaining with rabbit anti-PpRAD51 antibody (71) in 1:500 dilution along with DAPI on the squashed cells with protocol used for Arabidopsis root tips (70). Anti-rabbit secondary antibodies were conjugated with Alexa 488 (A27034; Thermo Fisher Scientific, Waltham, MA, USA) in 1:400 dilution in the Vectashield® mounting medium. Images were obtained with a Zeiss AxioObserver microscope and were analyzed using Zen Blue software. Images of plants, spores, and regenerating protoplasts were acquired with a Zeiss AxioZoom.V16 using Zen Blue software for acquisition and analysis. Detailed nuclei images of RAD51 foci and live imaging of GFP-BRCA2 germinating spore were taken using a LEICA SP5 II AOBS Tandem HyD confocal laser scanning microscope.

### Co-immunoprecipitation and yeast two-hybrid assays

The coimmunoprecipitation (CoIP) assays were performed on protein extract from the wild-type and GFP-BRCA2 protonema treated with or without 5 µg/ml bleomycin for 3h and ground in liquid nitrogen to a fine powder. Total cell lysates were prepared by adding lysis buffer (50 mM Tris pH 8.0, 150 mM NaCl, 0.1% Triton, 5% glycerol, 0.5% NP40, 0.5 mM EDTA, 1X protease inhibitor cocktail, 1X phosphatase inhibitor), with incubation on a rotating shaker for 30 min followed by centrifugation at 10 000 x g for 20 min at 4°C. For each CoIP, cell lysates corresponding to 5 mg total protein were incubated with GFP-Trap® magnetic particles M-270 (ChromoTek GmbH & Proteintech, Planegg-Martinsried, Germany) for 2 hours at 4°C. Protein-antibody complexes were recovered on magnetic particles after washing four times with lysis buffer. Protein complexes were then eluted with Laemmli buffer and migrated on SDS-PAGE. Proteins were detected by Western blot analysis with rat anti-GFP monoclonal antibody (cat no. 3h9, ChromoTek GmbH & Proteintech, Planegg-Martinsried, Germany) or anti-PpRAD51 antibody (71).

AH109 and Y187 (Clontech, TakaRabio, Shiga, Japan) yeast haploid strains were transformed with constructs encoding Gal4BD and Gal4AD fusion proteins for yeast two-hybrid assays. After mating the haploid yeast strains with opposite mating types on YPD plates, diploid yeast cells were selected on a dropout medium without leucine and tryptophan (SD/ -LW). Yeast-two-hybrid protein interactions were detected by plating five-fold serial dilutions (0 to 10, 000 fold) of each diploid on selective media depleted in leucine, tryptophan, histidine, and adenine (SD/ -LWH and SD/ -LWHA) for 3-5 days at 30°C. The strength of interaction was quantified by scoring each dilution that grew on the selective medium. Each dilution growing on SD/ -LWH or SD/ -LWHA was scored 0.05 or 0.1, respectively. The final score for each diploid was obtained by summing the individual scores, with 0.75 (0.25 + 0.5) being the maximum score for five dilutions growing on each selective medium without any self-activation.

### Protein expression and purification

The *P. patens* BRCA2 (PpBRCA2)-FL (amino acids 1 to 391), PpBRCA2-Cter (amino acids 251 to 391), PpBRCA2-Nter (1 to 250) proteins were expressed in *E. coli* BL21-Rosetta (DE3) (CamR). Expression clones containing recombinant pET15b-PpBRCA2 (AmpR) were grown at 37°C with chloramphenicol (25 µg/ml) and ampicillin (100 µg/ml) until the OD_600 nm_ reached 0.6. Recombinant protein expression was then induced by adding 0.2 mM IPTG followed by growth at 20°C overnight. After centrifugation (3000 rpm, 10 mins, 4°C), cells resuspended in lysis buffer (50 mM Tris–HCl pH 7.5, 300 mM NaCl, 5% glycerol, 40 mM imidazole, 1% triton X-100, 1 mM PMSF) were frozen in liquid nitrogen and lysed by sonication (Vibra-Cell™, VC 505) with proteases inhibitors (cOmplete™, EDTA-free Protease Inhibitor Cocktail - Roche).). Insoluble material was removed by centrifugation at (13, 000 rpm, 30 mins, 4°C). The clarified supernatant was purified by FPLC (Äkta pure, Cytiva).

For PpBRCA2-FL, PpBRCA2-Nter, and PpBRCA2-Cter purification, the clarified supernatant with 1 mM DTT was loaded onto a 5 mL HisTrap™ column (GE Healthcare) at a 2 mL/min flow rate using FPLC (Äkta pure, Cytiva). After washing with wash buffer (25 mM Tris-HCl pH 7.5, 500 mM NaCl, and 5% glycerol), proteins were eluted with an imidazole gradient (20 mL) with elution buffer (25 mM Tris-HCl pH 7.5, 500 mM NaCl, 5% glycerol, 500 mM imidazole). Fractions diluted to 50 mM NaCl were further purified via a 1 mL Resource Q column (Cytiva) using a linear gradient of 0.1 to 1 M NaCl. Fractions containing PpBRCA2-Nter or PpBRCA2-Cter were subjected to size exclusion chromatography on a Superdex® 75 10/300 GL column (Cytiva) equilibrated with 25 mM HEPES pH 7, 100 mM NaCl.

The *P. patens* DMC1 (PpDMC1) and RAD51 (PpRAD51-2) proteins fused to an N-terminal His6-SUMO tag, were expressed in *E. coli* BL21-Rosetta (DE3)-pLysS (CamR) using recombinant *pCDF-His6-Sumo-DMC1* and *pCDF-His6-Sumo-RAD51-2* by adding 0.2 mM IPTG. Cells were lysed in the lysis buffer (100 mM Tris–HCl pH 7.5, 3 M NaCl, 10% glycerol, 5 mM β-mercaptoethanol), and the cleared supernatant was loaded onto a 5 mL HisTrap™ column (GE Healthcare)) using FPLC (Äkta pure, Cytiva). After washing with wash buffer (25 mM Tris-HCl pH 8, 500 mM NaCl, 10% glycerol, 1 mM DTT, and 0.1 mM EDTA), proteins were eluted with an imidazole gradient (30 mL) in the wash buffer plus 400 mM imidazole. The SUMO-tag was cleaved by the SUMO protease (ratio of 200:1 (w:w)) overnight at 4°C during dialysis in 25 mM Tris-HCl pH 8, 500 mM NaCl, 10% glycerol, 5 mM β-mercaptoethanol. The sample was reloaded on a HisTrap™ column to recover the tag-free proteins in the flow through. The sample was further purified using a Hi-Trap™ heparin HP 5 ml column (GE Healthcare) with a 0.1-1M NaCl linear gradient. Fractions containing the protein of interest were pooled and diluted to reach 100 mM salt concentration and reloaded on a 6 mL Resource™ Q column (Cytiva) equilibrated with Tris-HCl pH 8, 100 mM NaCl, 10% glycerol, 1 mM DTT. Fractions containing PpDMC1 and PpRAD51-2 from elution with a 0.1-1M NaCl linear gradient were pooled, aliquoted, and stored at -80°C.

For NMR analysis, ^15^N-labeled proteins were produced in BL21-Rosetta (DE3) (CamR) grown in M9 medium containing 0.5 g/l ^15^NH_4_Cl and 2 g/l ^12^C-glucose.

### Nuclear Magnetic Resonance

2D NMR ^1^H-^15^N HMQC experiments were conducted on a 3-mm sample tube containing ^15^N-labeled PpBRCA2, PpBRCA2-Nter, and PpBRCA2-Cter at 20µM, 200µM and 260µM concentrations, respectively in 25 mM HEPES pH 7.0, 100 mM NaCl, with or without 1 mM DTT, using a 950 MHz Bruker Advance III spectrometer with a triple resonance cryogenic TCI probe at 280 K.

### BioLayer Interferometry (BLI)

Biomolecular interactions between recombinant recombinases (PpRAD51-2, PpDMC1) and PpBRCA2-derived peptides were analyzed by BLI using an Octet RED96 instrument (FortéBio). Biotinylated PpBRCA2 peptides were synthesized by ProteoGenix, France. Peptide sequences are given in DATASET_S3.

Briefly, all BLI measurements were conducted at 25 °C. The biosensors were first hydrated for 10 min in buffer A (25 mM Tris, pH 7.5, 100 mM Na_2_SO_4_, 5 mM β-mercaptoethanol) or buffer B (25 mM Tris, pH 7.5, 100 mM NaCl, 5 mM β-mercaptoethanol). The biotinylated peptides were immobilized by incubating Streptavidin (SA) biosensors in 0.5 µM peptide solutions. For the binding assays, the buffers used were supplemented up to 0.05% Tween-20. To monitor the association, prewashed biosensors were incubated with PpRAD51-2 for 300 sec. Kinetic assays were performed at varying concentrations of PpRAD51-2 using two-fold serial dilutions (except for the peptide BRC3G). For PpDMC1, SA biosensors were incubated with 10 µM protein for 300 sec. The dissociation was followed for 600 seconds and the dissociation constants (Kd) were determined in steady-state mode. After each association and dissociation round, biosensors underwent five regeneration cycles (10 sec in 1 M NaCl or 5 M NaCl for BRC3G peptide, followed by 10 sec in buffer A or B with 0.05% Tween-20).

For interactions between PpBRCA2-FL, PpBRCA2-Nter, PpBRCA2-Cter, and biotinylated single-stranded DNA (100 nucleotides) by BLI, the SA biosensors were hydrated for 10 min in 25 mM Tris, pH 7.5, 100 mM NaCl or 25 mM HEPES, pH 7, 100 mM NaCl, 1 mM DTT buffer. The biotinylated DNA (100 nucleotides (nt) ssDNA) was immobilized on SA biosensors using a 50 nM solution. For the binding assays, the buffers used were supplemented up to 0.05% Tween-20. After washing, sensors were incubated with PpBRCA2-FL, PpBRCA2-Nter, and PpBRCA2-Cter at varying concentrations with two-fold serial dilutions for 300 sec to monitor association, followed by 600-sec dissociation in a buffer.

Biomolecular interactions between PpRAD51-2 and 6His-PpBRCA2-Cter were measured using Ni-NTA biosensors (NTA), prehydrated for 10 min in 25 mM Tris, pH 7.5, 100 mM NaCl, 1 mM DTT buffer. The 6His-PpBRCA2-Cter protein was immobilized on NTA sensors through sensor immersion in 0.5 µM solution. For the binding assays, the buffer 25 mM Tris, pH 7.5, 100 mM Na_2_SO_4_, 5 mM β-mercaptoethanol, 0.1% Tween-20 was used. After washing, sensors were incubated with PpRAD51-2 at varying concentrations with two-fold serial dilutions for 300 sec to monitor association, followed by 600-sec dissociation in a buffer.

### Electromobility mobility shift assay (EMSA)

For PpBRCA2-DNA binding assays, 2 µM nucleotides (nt) (20 nM molecules) of a ssDNA (100 nt), dsDNA (250 bp) or a ss-ds overhang (150 nt - 100 bp) Cy5 labeled DNA substrate were incubated with PpBRCA2-FL, PpBRCA2-Cter or PpBRCA2-Nter (0 - 2.4 µM), 7 min at 37°C in a buffer containing 10 mM Tris-HCl pH 7.5, 50 mM NaCl, 5 MgCl_2_, 2 mM ATP and 1 mM DTT. Protein–DNA complexes were fixed by adding 0.01 % glutaraldehyde 5 min at room temperature. The reaction products were run onto a 0.75 % agarose gel in 0.5 X Tris-acetate-EDTA (TAE) at 80 V and 4°C for 20 min. The gel was scanned on a Typhoon imager (GE Healthcare Life Science) using the Cy5 channel. The percentage of DNA-protein complexes was estimated in each condition by Image J software measuring the intensity of the displaced bands (corresponding to the DNA-protein complexes) normalized to the total intensity of all bands (corresponding to free DNA plus DNA-protein complexes).

To test the interaction of ssDNA-PpRAD51-2 filaments with PpBRCA2-Cter or BRC4, 3 µM nucleotides (nt) (30 nM molecules) of a ssDNA (100 nt), Cy5 labeled DNA substrate were incubated with PpRAD51-2 and PpBRCA2-Cter (0 - 4 µM) or PpBRCA2-BRC4 (0 – 4 µM), 15 min at 37°C in a buffer containing 10 mM Tris-HCl pH 7.5, 50 mM NaCl, 5 MgCl_2_, 2 mM ATP and 1 mM DTT. Protein– DNA complexes were fixed by adding 0.01 % glutaraldehyde 5 min at room temperature. The reaction products were run onto a 0.75 % agarose gel in 0.5 X TAE at 80 V and 4°C for 20 min. The gel was scanned on a ChemiDoc imager (BIO-RAD) using the Cy5 channel.

### AlphaFold3 predictions

AlphaFold3 predictions for protein complexes were computed through Alphafold3 online tool (https://alphafoldserver.com/welcome). The plDDT, and ipTM scores and graphs were provided directly by the program. Molecular graphics and analyses were performed with the UCSF ChimeraX (version 1.7.3), developed by the Resource for Biocomputing, Visualization, and Informatics at the University of California, San Francisco.

## RESULTS

### 1. Identification of a non-canonical BRCA2 homolog in *Physcomitrium patens*

BRCA2 homologs are present in diverse plants but were not identified in *Physcomitrium patens* (52, 33), a moss model species from the bryophyte clade in the Plantae kingdom. Among plants, two *Arabidopsis thaliana* AtBRCA2A and AtBRCA2B homologs possess three key features of eukaryotic BRCA2 orthologs: BRC repeats, a canonical DBD, and a putative nuclear localization signal (NLS) (9, 33, 72). We performed PSI-BLAST searches with AtBRCA2A as a query but failed to identify any homolog in *P. patens.* We employed a more sensitive sequence search method, HHpred, to detect remote homology based on the comparison of profile hidden Markov models (62). Using AtBRCA2A as bait and targeting HHpred searches to human, drosophila, and moss proteomes with the default PDB_mmCIF70_16_Aug database, we retrieved 147 hits including human and drosophila BRCA2 homologs as well as other DBD-containing proteins such as RPA (DATASET_S1). Among the *P. patens* hits, proteins without BRC repeats were ruled out to be BRCA2 homologs. We detected only one sequence corresponding to the protein XP_001764308.1/Pp6C10_10830 with a probability of 94.7% and a pairwise sequence identity of 31% with the N-terminal region of AtBRCA2A (first 506 amino acids). The HHpred pairwise alignment revealed the presence of BRC repeats in Pp6C10_10830 (Fig. S1A). Conversely, HHpred search using Pp6C10_10830 retrieved BRCA2 homologs from Arabidopsis, human, and drosophila with pairwise alignments showing strong homology in the BRC repeat region, substantiating that Pp6C10_10830 is homologous to other BRCA2 proteins (DATASET_S2). The *P. patens Pp6C10_10830* gene consists of a single exon without any intron located on chromosome 10 and encodes a 391 amino acids (aa) long protein, which we designated as PpBRCA2 (Fig. S1B). PpBRCA2 possesses a nuclear localization signal (NLS) and four BRC repeats in its C-terminal region (Fig. 1A, Fig. S1A&B). The alignment of BRC repeats detected two fully conserved FxxA and dihydrophobic motifs across the moss, *A. thaliana*, human, and drosophila (Fig. 1B), corresponding to previously reported two modules of BRC repeats (39, 35). Alphafold3 (AF (73)) predicted interactions with strong confidence scores (ipTM > 0.7) between moss BRC repeats 1, 2, and 4 and both moss RAD51 homologs (Fig. 1C, Fig. S1D). The superimposition of calculated models and the crystal structure available for human BRC4 bound to RAD51 revealed that the conserved phenylalanine residue of the FxxA motif is buried within the RAD51 ATPase domain (Fig. S1E; (38)). The BRC3-RAD51 interaction scores were lower (ipTM < 0.7), with the phenylalanine of the FxxA motif not positioned in the ATPase domain of RAD51 (Fig. 1C, Fig. S1D). These AF results are consistent with previous reports showing differences in interaction affinities of BRC repeats towards RAD51 (37, 74, 34).

**Figure 1.**
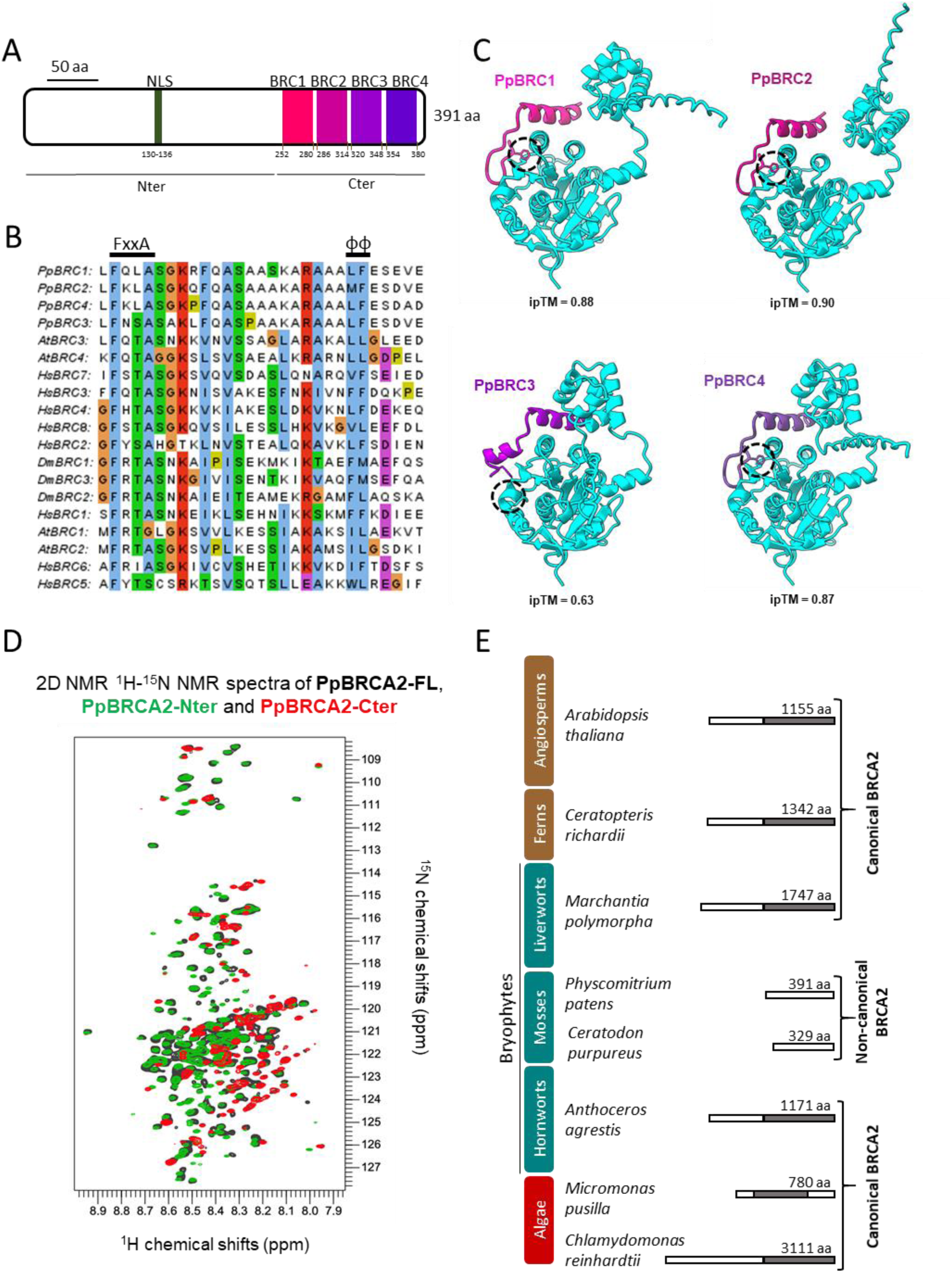
Identification of non-canonical BRCA2 homolog in *Physcomitrium patens*. A. Schematic representation of *P. patens* BRCA2 (PpBRCA2) with predicted nuclear localization signal (NLS) in green and four conserved BRC repeats in pink, orchid, violet and purple. B. Alignment of BRC repeats from *Homo sapiens (Hs), Arabidopsis thaliana (At), Drosophila melanogaster (Dm)* and *Physcomitrium patens (Pp)*. Conserved FxxA and dihydrophobic (φφ) motifs are labeled with amino acid residues in Clustal X color code. C. 3D structure models of the interaction between PpRAD51-1 (cyan) and PpBRCA2 BRC1(pink), BRC2 (orchid), BRC3 (violet), BRC4 (purple) built with AlphaFold3. The phenylalanine of the conserved FxxA motif in BRC repeats is shown in dotted circles. The ipTM scores assessing the accuracy of the predicted interfaces are presented under each model. D. Superimposition of the 2D NMR ^1^H-^15^N SO-FAST HMQC spectra of PpBRCA2 (black), PpBRCA2-Nter (green), and PpBRCA2-Cter (red) recorded at 950 MHz, showing that all constructs are intrinsically disordered. E. Evolutionary conservation of canonical BRCA2 homologs with DNA binding domain (DBD) in grey or non-canonical BRCA2 without DBD in Algae (Red), Bryophytes (Cyan) and Tracheophytes (Brown) of the Plantae kingdom.

Strikingly, the HHpred pairwise alignment revealed that PpBRCA2 lacks a canonical DBD, which typically includes a well-folded helical domain and three OB domains (Fig. 1A, Fig. S1B &C). We purified recombinant full-length PpBRCA2, PpBRCA2-Nter (aa 1-250 aa), and PpBRCA2-Cter (251-391 aa) to determine the presence of folded domains by nuclear magnetic resonance (NMR) spectroscopy. Our NMR analysis confirmed that recombinant PpBRCA2, PpBRCA2-Nter and PpBRCA2-Cter are highly disordered in solution (Fig. 1D), except the repeated BRC sequences giving rise to stretches of partially overlapping peaks at the C-terminal (Fig. 1D). Our NMR data indicated the absence of the well-folded domain, suggesting a lack of folded DBD in PpBRCA2. Thus, PpBRCA2 might not require a folded DBD for its functions like DmBRCA2, or an unknown protein might compensate for the loss of canonical DBD. Among the well-annotated moss genomes, PpBRCA2 homologs without DBD were identified using PSI-BLAST searches in *Ceratodon purpureus* (CepurR40.9G141900 or CepurGG1.9G149800) and *Funaria hygrometrica* (Fh_23015, (75)) species, indicating that loss of DBD is common to at least three moss genera (Fig. 1E, Fig. S1C). Since establishing phylogenetic relationships among BRCA2 orthologs is challenging due to varying protein sizes and low aa conservation outside the BRC repeats and DBD (41), we compared the presence or absence of the canonical DBD across various plant clades. BRCA2 homologs with canonical DBD were readily detected in the clades of green algae, ferns, and angiosperms, as well as in both liverworts and hornworts – which together with mosses form the bryophyte group, suggesting that BRCA2 homologs without a canonical DBD have likely emerged in moss lineages only (Fig. 1E). Overall, two variants of BRCA2 orthologs appear to exist: canonical BRCA2 with a canonical DBD and non-canonical BRCA2 homologs lacking this DBD (Fig. S1F).

### 2. PpBRCA2 is essential for normal plant growth and DSB repair during meiosis

The life cycle of *P. patens* involves haploid spores developing into filamentous protonema, leading to gametophores on leafy shoots, which produce diploid sporophytes upon fertilization. Meiosis occurs in the sporophyte and generates haploid spores to complete the life cycle. We explored the function of PpBRCA2 by analyzing four deletion mutants generated using CRISPR-Cas9 genome editing in Reute (*brca2Δ-1* and *brca2Δ-2*) and Gransden (*brca2Δ-3* and *brca2Δ-4*) *P. patens* ecotypes (Fig. S2A). All four *brca2Δ* mutants were viable but exhibited a significant reduction (∼30%) in plant diameter compared with wild-type (WT) (Dunnett test P<0, 0001), indicating a slower protonema growth in the absence of PpBRCA2 (Fig. 2A&B, Fig. S2C). Both WT and *brca2Δ* mutants developed sporophyte and produced spores; however, *brca2Δ* mutant spores were irregularly shaped, heterogeneous in size, and generally smaller than the round, larger WT spores (Fig. 2A, Fig. S2F). Spore germination test showed that only 2% of *brca2Δ* spores germinated compared with 70% of WT spores, suggesting that *brca2Δ* mutants predominantly produce non-viable spores (Fig. 2C, Fig. S2D&E). The few germinated *brca2Δ* spores displayed abnormal protonema with severely reduced growth, potentially due to aneuploidy (Fig. S2B). Thus, PpBRCA2 is essential for normal development and spore viability.

**Figure 2.**
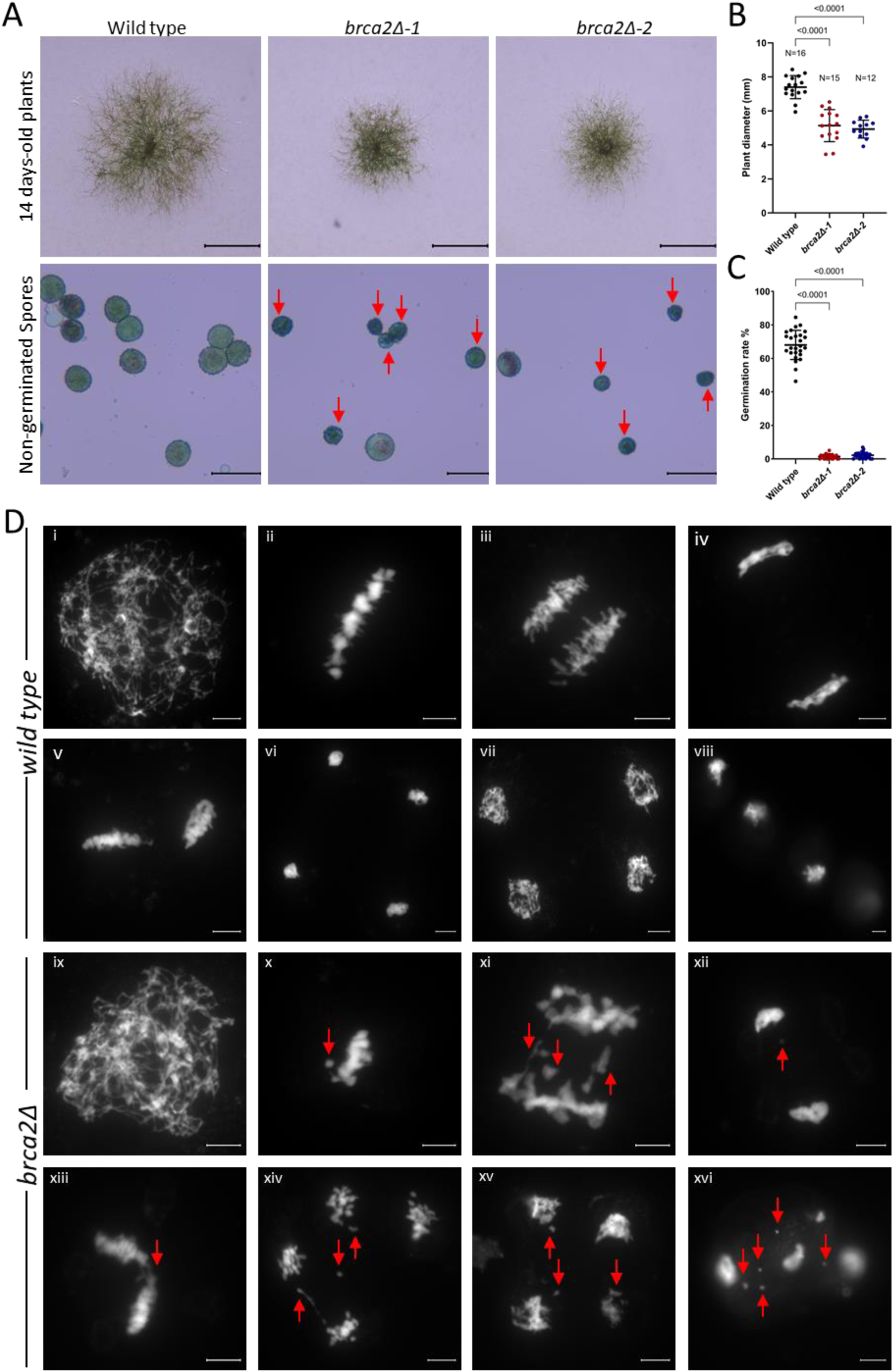
*P. patens brca2Δ* mutants display growth retardation, non-viable spores, and meiosis defects. A. Top row: representative images of 14-day-old wild-type and *brca2Δ* plants on minimal media. Scale bars: 1 mm. Bottom row: images of wild-type and *brca2Δ* non-germinated spores stained with Alexander stain. Red arrows point to an anomaly in the shape and size of mutant spores. Scale bars:50 µm. B. Quantification of 14-day-old plant diameter on minimal media. Each dot represents the diameter of an individual plant for wild type, *brca2Δ-1*, and *brca2Δ-2* genotypes. Horizontal bars indicate the mean and standard deviation. The p-values shown were calculated using one-way ANOVA with Dunnett’s test for multi-comparison. C. Quantification of spore germination rate after 6 days of growth. Spores were pooled from four different capsules for each genotype. Each dot represents the germination percentage of spots containing 8 to 25 spores, with total spore counts in wild type = 3357, *brca2Δ-1* = 2093, and *brca2Δ-2* = 1715. Horizontal bars indicate the mean and standard deviation. The p-values shown were calculated using one-way ANOVA with Dunnett’s test for multi-comparison. D. Representative images of DAPI-stained chromosome spreads at leptotene (i, ix), metaphase I (ii, x), anaphase I (iii, xi), telophase I (iv, xii), metaphase II (v, xiii), anaphase II (xiv), telophase II (vi, vii, xiv), and mature spores (viii, xvi) in wild type (i-viii) and *brca2Δ* (ix-xvi). Red arrows indicate lagging chromosomes, DNA fragmentation, or univalent chromosomes. Scale bars: 5 µm.

*P. patens* haploid spores are the direct product of the meiosis in the sporophytes, suggesting that non-viable *brca2Δ* spores may result from defective meiosis (58). To examine this, we analyzed the meiotic progression in WT, *brca2Δ-1*, and *brca2Δ-2* sporophytes by preparing meiotic chromosome spreads with DAPI staining. In WT, meiotic chromosomes undergo DSB formation and HDR-mediated repair during prophase I, culminating in bivalent formation at metaphase I, where homologous chromosomes are held together by crossovers (Fig. 2D). These bivalents ensure the balanced segregation of homologous chromosomes at anaphase I (Fig. 2D). During the second meiotic division, sister chromatids align at metaphase II and separate at anaphase II (Fig. 2D). Following telophase II and cytokinesis, four haploid cells (tetrad) form, completing meiosis and leading to spore production (Fig. 2D). We observed that 95% of metaphase I cells (N=41) displayed only bivalents, while <5% of cells (N=2) showed a mixture of bivalents and univalents (chromosomes without crossover). In *brca2Δ* mutants, we detected a significantly high frequency of aberrant metaphase I cells (21%, N=12 out of 56) in which some DAPI-stained chromatin remained outside the aligned bivalents (Fig. 2D). Due to high chromatin compaction and low resolution, we were unable to determine whether these signals represented univalents or fragmented chromosomes. However, we consistently observed chromosome bridges and micronuclei in later meiotic stages and mature spores (N=71) (Fig. 2D), indicating severe meiotic defects. These abnormalities suggest a failure in meiotic break repair, leading to unbalanced chromosome segregation in *brca2Δ* mutants, similar to defects observed in *rad51-1-2* mutant meiocytes (Fig. S3). Taken together, PpBRCA2 is required for DSB repair during meiosis in *P. patens*.

### 3. PpBRCA2 is essential for somatic DNA repair and genome stability

To determine the role of PpBRCA2 in somatic DNA repair, we analyzed the sensitivity of *brca2Δ* mutants to DNA-damaging agents. We exposed WT and *brca2Δ* (*brca2Δ-1, -2, -3,* and *-4*) regenerating protoplasts to various dosages of UV-B, which induces pyrimidine dimers producing single- and double-strand breaks during replication (76). After a one-week recovery period, *brca2Δ* mutants exhibited a significantly lower survival rate than WT at 60 and 120 mJ/cm², indicating that *brca2Δ* mutants are hypersensitive to UV-B (Fig. 3A, Fig. S4A, C&E). Next, we tested the response of *brca2Δ* mutants to bleomycin, a drug that induces DSBs (77). WT and *brca2Δ* protoplasts were plated on media supplemented with increasing concentrations of bleomycin, and cell regeneration was assessed after 72 hours. *brca2Δ* mutants exhibited a significant reduction in regeneration frequency at 1.6, 8, and 40 ng/ml compared with WT, further supporting a role for PpBRCA2 in DSB repair (Fig. 3B, Fig. S4B, D&F). These findings demonstrate that PpBRCA2 is essential for DNA repair and genome stability in *P. patens*.

**Figure 3.**
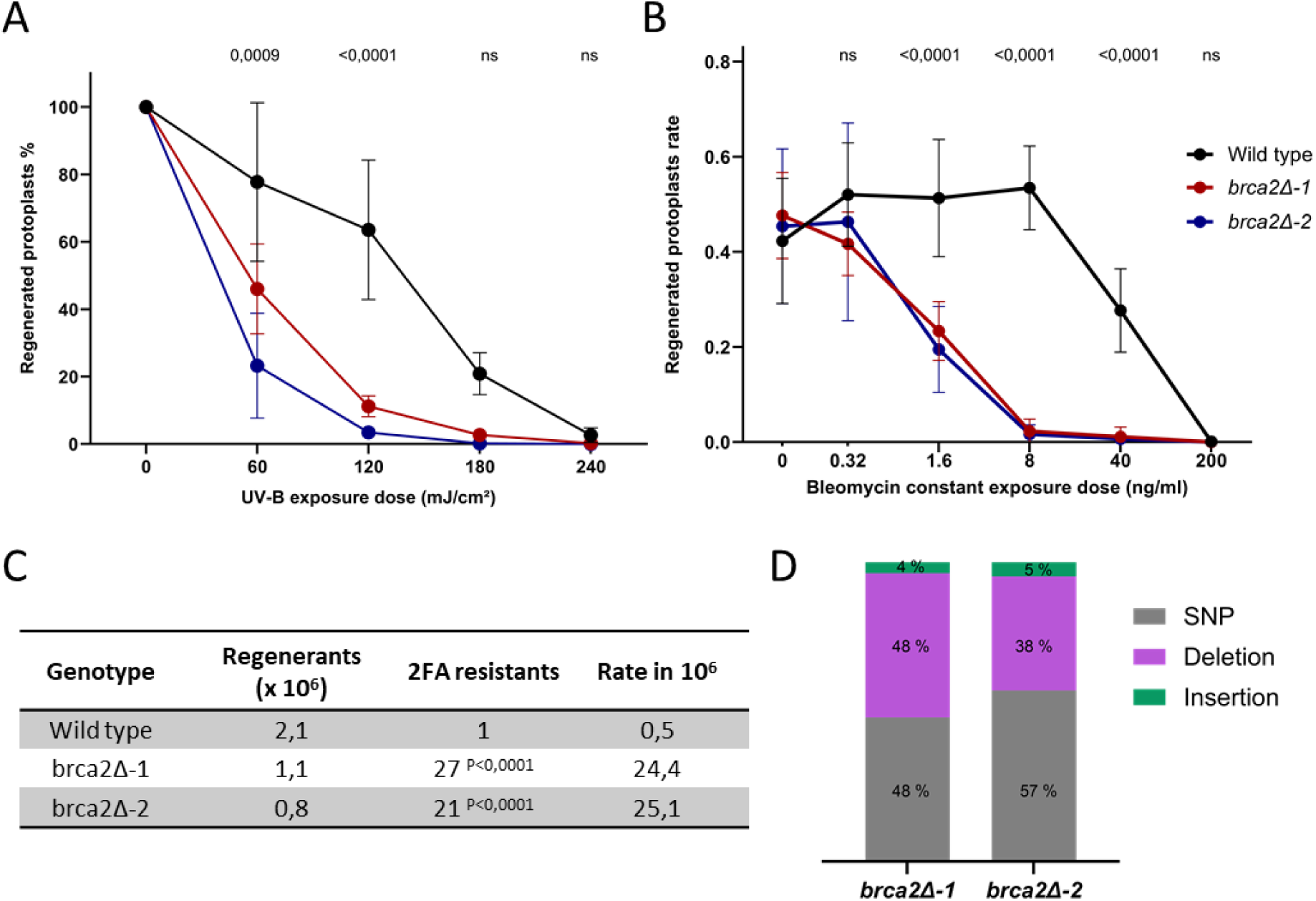
*P. patens brca2Δ* mutants display growth retardation, non-viable spores, and meiosis defects. A. Survival curves of wild-type (black), *brca2Δ-1* (red) and *brca2Δ-2* (blue) protoplasts regenerating 6 days post-exposure to UV-B. Values are normalized on the non-treated sample. For each point, the mean of three independent experiments is plotted with error bars indicating standard deviation. The p-values were calculated using two-way ANOVA followed by Dunnett’s test for multi-comparison. B. Survival curves of wild-type (black), *brca2Δ-1* (red) and *brca2Δ-2* (blue) protoplasts regenerating after 3 days of constant exposure to bleomycin. Values are independent. For each point, the mean of three independent experiments is plotted with error bars indicating standard deviation. The p-values shown were calculated using two-way ANOVA followed by Dunnett’s test for multi-comparison. C. Rate of spontaneous mutations at the *APT* locus in wild-type and *brca2Δ* regenerating protoplasts. The following p-values were calculated using the Fisher exact test. D. Histograms show the distribution of single nucleotide polymorphism (SNP), deletion or insertions identified by sequencing in *brca2Δ-1* and *brca2Δ-2* at the *APT* locus.

We next evaluated whether the loss of PpBRCA2 leads to genetic instability by measuring the spontaneous mutation rate at the *APT* (adenine phosphoribosyl transferase) reporter gene as previously described (78). The 2-fluoroadenine (2FA) is a toxic adenine analog for WT and allows the selection of plants harboring nonsense mutations in the *APT* gene. WT and *brca2Δ* (*brca2Δ-1* and *brca2Δ-2*) protoplasts were plated on media containing 10 µM 2FA to obtain 2FA-resistant plants. A total of 2.1 x 10^6^, 1.1 x10^6^, and 0.8 x 10^6^ protoplasts resulted in 1, 27, and 21 2FA-resistant plants in WT, *brca2Δ-1,* and *brca2Δ-2*, respectively. This corresponded to a spontaneous mutation rate of 0.5 x 10^6^ in WT, 24.4 x 10^6^ in *brca2Δ-1,* and 25.1 x 10^6^ in *brca2Δ-*2, indicating a 50-fold increase in mutation accumulation rate in *brca2Δ* mutants compared with WT (Fig. 3C). Sequencing of the *APT* locus in 2FA-resistant *brca2Δ* plants revealed about half of the mutations were single nucleotide polymorphisms (SNPs), leading to amino acid changes with premature stop codons, or splicing site alterations (Fig. 3D, DATASET_S3). Rest of mutations consisted of deletions ranging from 1 to 283 bp, while < 5% of resistant plants showed insertions, mostly 1-2 bp except one being 37 bp insertion. Notably, all insertions except one were accompanied by deletions, often displaying 2–4 bp microhomology at the repair junctions. The only WT-resistant plant carried an SNP affecting a splice site at the exon 3 border (Fig. 3D, DATASET_S3). These findings support that PpBRCA2 plays a crucial role in repairing naturally occurring DNA damage and in preventing genome-wide mutation accumulation, a conserved function of BRCA2 homologs (6).

### 4. Loss of PpBRCA2 disrupts somatic HDR and gene targeting

Given the conserved role of BRCA2 orthologs in HDR, we assessed homology-dependent gene targeting (GT) efficiency at the *APT* locus in WT and *P. patens brca2* mutants. For GT assays, we co-transfected protoplasts with three plasmids: one expressing the Cas9 gene, the second expressing the sgRNA#2 targeting the *APT* gene, and the third carrying a donor DNA cassette with a G418-resistant gene flanked by the APT gene sequences homology arms (Fig. S5) (69). In our GT assays, the DSB repair at the *APT* locus can result in three genotypes: (1) 2FA-resistant and G418-sensitive plants – arising from non-HDR (e.g. classical NHEJ or alternative end-joining (Alt-EJ)) repair of the DSBs that can lead to mutations but without cassette integration. (2) 2FA- and G418-resistant plants - originating from HDR-mediated GT with the donor DNA cassette integration at the *APT* locus. (3) 2FA- and G418-resistant plants - derived from non-HDR repair of *APT* locus breaks, with illegitimate donor cassette integration elsewhere in the genome. Following the protoplast transfection, plants were selected on 2FA and tested for G418 resistance. The majority of plants (83.5 % in WT and 98.4% in *brca2Δ* mutants) were 2FA-resistant with no G418 cassette integration, likely resulting from NHEJ or Alt-EJ repair (Fig. 4A), suggesting *PpBRCA2* is not essential for non-HDR pathways. However, the GT rate -measured as the frequency of 2FA- and G418-resistant plants - was ∼10-fold lower in *brca2Δ* mutants (1.6%) than in WT (16.5%), similar to *rad51-1-2* (69). This demonstrates that PpBRCA2 is essential for HDR-mediated GT.

**Figure 4.**
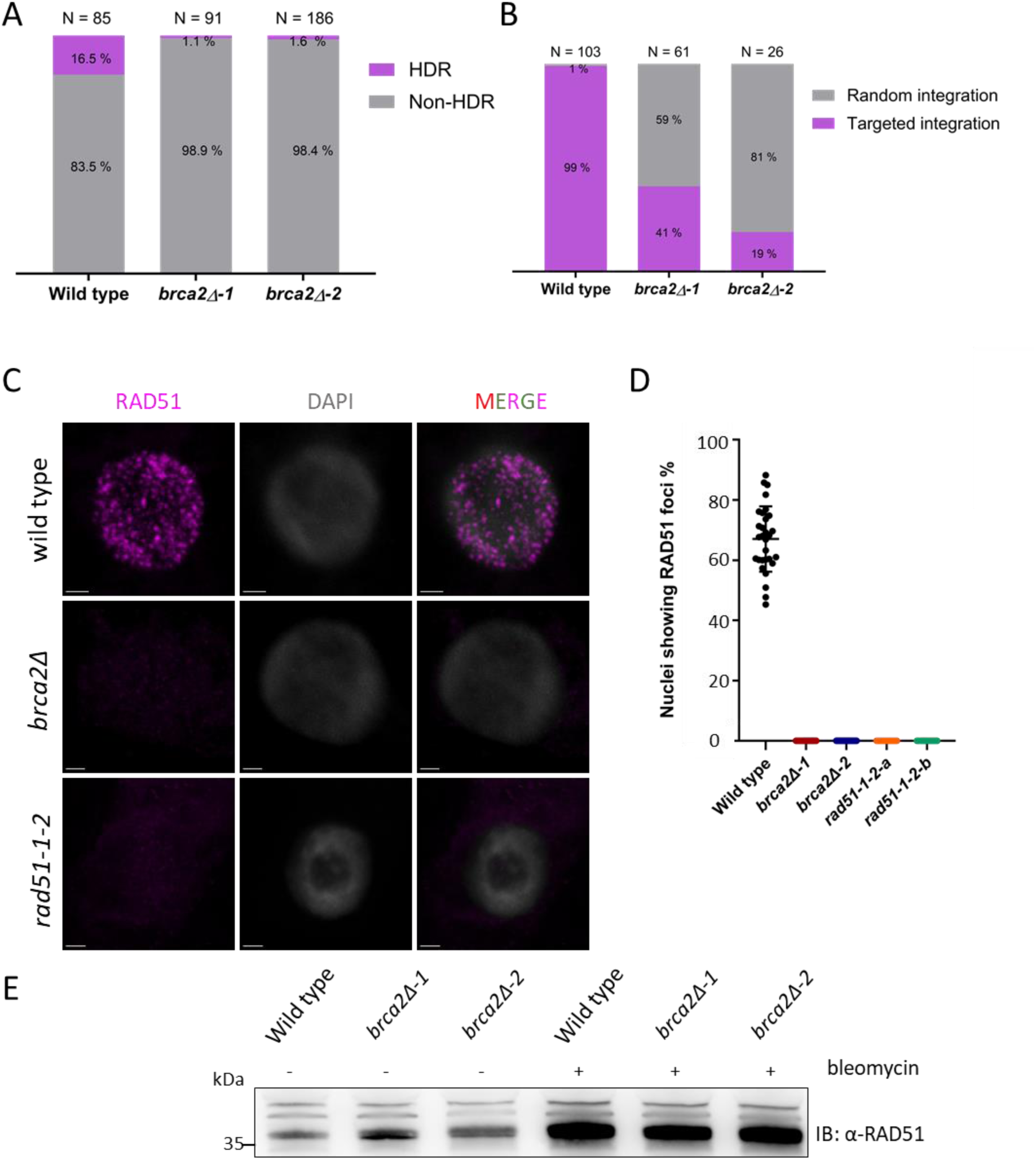
Analysis of homologous recombination repair (HDR) and RAD51 recruitment in *P. patens* wild-type and *brca2Δ* mutants. A. Percentage of gene targeting events (donor G418 cassette insertion) by HDR (purple) and non-HDR events (no insertion) in grey at the *APT* locus in wild-type, *brca2Δ-1,* and *brca2Δ-2* protoplasts. Cassette integrations were confirmed by genotyping G418^R^ and 2-FA^R^ plants using *APT*-locus-specific primers. B. Percentage of targeted (purple) and random integration (grey) of the donor G418 cassette at the *APT* locus in wild-type and *brca2Δ* protoplasts. Cassette integrations were confirmed by genotyping stable G418^R^ plants using *APT*-locus-specific primers. C. Immunolocalization of RAD51 (purple) and DNA marked with DAPI (grey) on squashed protonema cells from wild-type, *brca2Δ,* and *rad51-1-2* plants treated with 5 µg/ml bleomycin for 3 h. Scale bars: 2 µm. D. Percentage of RAD51 foci positive cells from wild-type, *brca2Δ,* and *rad51-1-2* plants. Each dot represents the percentage of RAD51 foci positive nuclei from various images with total cells counted in wild type = 1698, *brca2Δ-1* = 1083, *brca2Δ-2* = 1554, *rad51-1-2-a* = 385 and *rad51-1-2-b* = 404. Horizontal bars indicate the mean and standard deviation. E. Western blot analysis of RAD51 steady-state levels from wild type and *brca2Δ* moss protonema total protein extract with or without 3 hours treatment with 5 µg/ml bleomycin probed with anti-RAD51 antibody.

Since HDR impairment enhances random DNA integration in *P. patens* (58, 69), so we also examined the frequency of targeted vs. random integration of the donor cassette by selecting WT and *brca2Δ* mutants on G418 and then testing for 2FA resistance. Targeted integration is confirmed by the PCR-detected cassette insertion at the *APT* locus, while random integrations lacked the *APT* locus integration in stable G418-resistant plants (Fig. S5). In WT, 99% of G418-resistant plants had targeted integration events, with only 1% showing random integration. In contrast, *brca2Δ* plants exhibited a severe reduction in targeted integration events (<41%), indicating a higher rate of random insertions (Fig. 4B).

Our data argue that PpBRCA2 promotes HDR to counteract illegitimate integration of exogenous DNA and reinforce the importance of PpBRCA2 for genome stability in *P. patens*.

### 5. PpBRCA2 is essential to RAD51 focus formation

After observing defective HDR in *brca2Δ* mutants, we examined whether PpBRCA2 mediates the recruitment of RAD51, detectable as nuclear foci following DSB induction. Protonema filaments treated with or without 5 µg/ml of bleomycin for 3 hours were immunostained with anti-RAD51 antibody and DAPI. No RAD51 foci were detected in untreated WT and *brca2Δ* cells (Fig. S3E). Immunostaining of *rad51-1-2* mutants served as a control of anti-RAD51 antibody specificity, showing no RAD51 signal (Fig. 4C). After bleomycin treatment, numerous RAD51 nuclear foci were observed in ∼70% of protonema cells in WT (Fig. 4C&D). In contrast, no *brca2Δ* protonema cells showed RAD51 foci after bleomycin treatment (Fig. 4C&D). The absence of RAD51 foci in *brca2Δ* mutants could be due to low RAD51 protein levels. However, our Western blot analysis revealed similar RAD51 protein abundance in WT, *brca2Δ-1,* and *brca2Δ-2* mutants treated with bleomycin (Fig. 4E), suggesting that despite producing WT levels of RAD51, *brca2Δ* mutants fail to load RAD51 at DSB sites. These results show that similar to other BRCA2 orthologs, PpBRCA2 is essential for RAD51 focus formation after DSB formation.

### 6. A non-canonical *Ceratodon purpureus* BRCA2 can complement *P. patens brca2Δ* mutant

We identified BRCA2 homologs in three moss species without DBD, though the number of BRC repeats varied among them. In contrast to PpBRCA2, which contains four BRC repeats, *F. hygrometrica* BRCA2 and *C. purpureus* BRCA2 (CpBRCA2) have three and one BRC repeat(s), respectively. We further explored whether a non-canonical BRCA2 with only one BRC repeats could complement the functions of PpBRCA2. Additionally, we examined whether AtBRCA2A without DBD (AtBRCA2A^ΔDBD^) could functionally replace PpBRCA2. We integrated the coding sequence for either *CpBRCA2* or *AtBRCA2A^ΔDBD^*omitting the canonical DBD, at the *PpBRCA2* locus in *brca2Δ-1*. As a control, we integrated the *PpBRCA2* gene carrying 15 silent polymorphisms (designated as *PpBRCA2*),* which distinguished it from the WT *PpBRCA2* sequence (Fig. S6). We isolated independent lines with the integration of each *BRCA2* gene at the *PpBRCA2* locus by genotyping and sequencing. Each selected line was evaluated for restoration of the growth phenotype of *brca2Δ-1*. The integration of *PpBRCA2\** restored a WT level of plant growth in *brca2Δ-1* (Fig. 5A&B). The *CpBRCA2* integrated plants showed about 10% reduction in plant diameter compared with WT (Fig. 5A&B), suggesting CpBRCA2 can largely rescue the growth phenotypes of *brca2Δ-1*. *AtBRCA2A^ΔDBD^* integrated plants, in contrast, showed a severe reduction in diameter compared with WT (Fig. 5A&B), but similar to *brca2Δ-1*, suggesting that *AtBRCA2A^ΔDBD^* failed to complement *brca2Δ-1* mutant. To examine in vivo the mediator role of PpBRCA2*, CpBRCA2, and AtBRCA2A^ΔDBD^, we performed immunolocalization of RAD51 on protonema cells treated with bleomycin from the three corresponding genotypes and WT. We detected RAD51 foci in >90% of cells in WT and protonema cells complemented with either *PpBRCA2\** or *CpBRCA2*, while *AtBRCA2^ΔDBD^* protonema showed no RAD51 foci, except a diffuse nuclear signal in some cells (Fig. 5C&D, Fig. S6B). This diffuse signal in the *AtBRCA2^ΔDBD^* line may reflect the accumulation of RAD51 in the nucleus. Western blot analysis confirmed the abundance of RAD51 at the WT level in response to DSB induction in PpBRCA2*, CpBRCA2, and AtBRCA2^ΔDBD^ plants, suggesting a likely defect in RAD51 recruitment in *AtBRCA2^ΔDBD^* (Fig. S6C). Using RT-PCR analysis, we confirmed the expression of *PpBRCA2\**, *CpBRCA2*, and *AtBRCA2^ΔDBD^* by detecting their respective transcripts (Fig. S6D). These findings together demonstrate that CpBRCA2 with only a single BRC repeat is sufficient to support PpBRCA2 functions during HDR.

**Figure 5.**
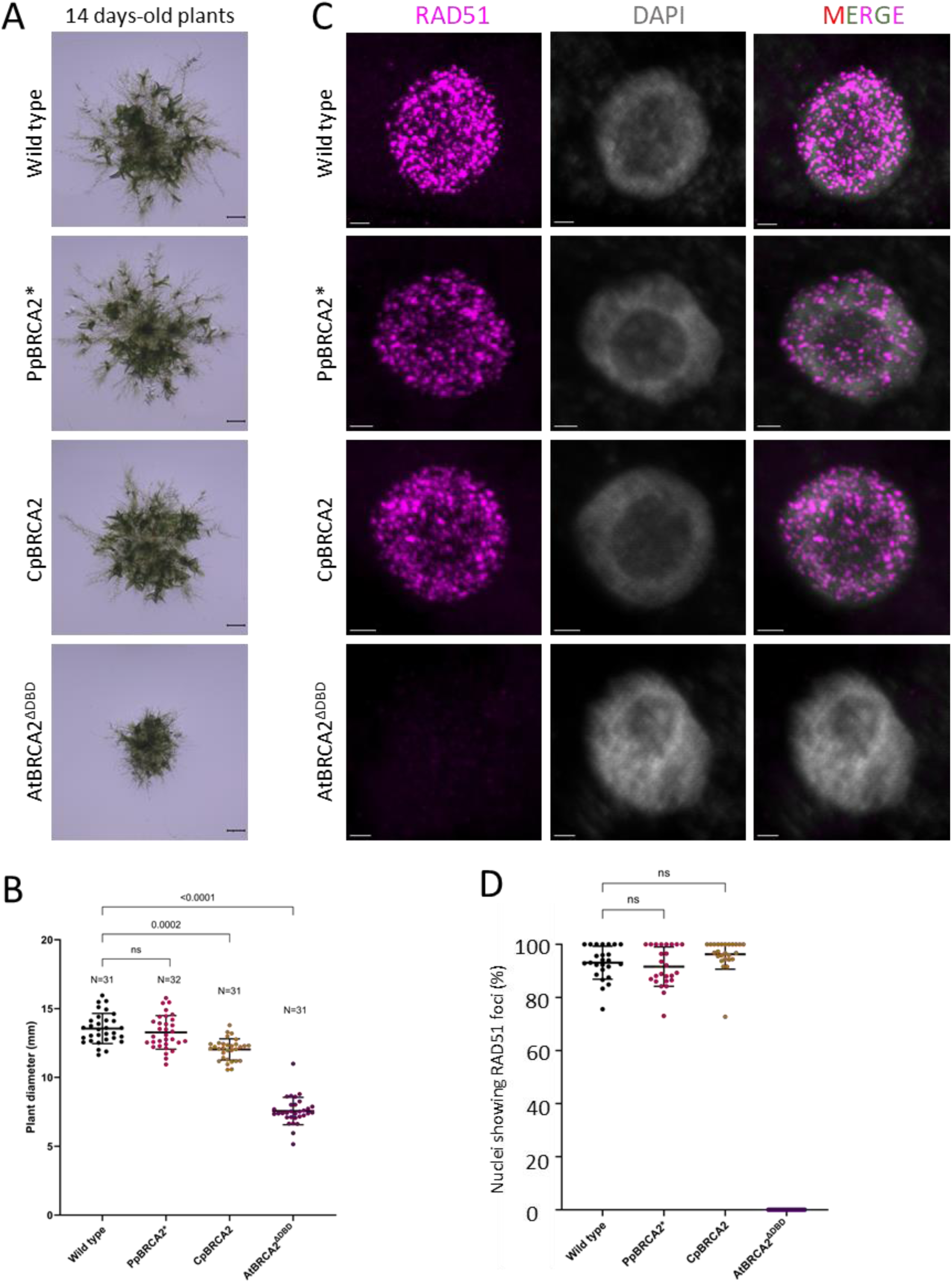
Growth phenotype and RAD51 recruitment in *P. patens brca2Δ-1* complemented plants. A. Representative images of 14-day-old wild type, PpBRCA2* (*brca2Δ-1)*, CpBRCA2 (*brca2Δ-1*) and AtBRCA2^ΔDBD^ (*brca2Δ-1*) plants. Scale bars: 1 mm. B. Quantification of 14-day-old plant sizes on minimal media. Each dot represents the diameter of an individual plant for wild type, PpBRCA2* (*brca2Δ-1)*, CpBRCA2 (*brca2Δ-1*), and AtBRCA2^ΔDBD^ (*brca2Δ-1*) genotypes. Horizontal bars indicate the mean and standard deviation. The p-values shown were calculated using one-way ANOVA with Dunnett’s test for multi-comparison, ns; non-significant. C. Immunolocalization of RAD51 (purple) and DAPI staining (grey) on the squashed protonema cells of wild type, PpBRCA2* (*brca2Δ-1)*, CpBRCA2 (*brca2Δ-1*) and AtBRCA2^ΔDBD^ (*brca2Δ-1*) plants treated with 5 µg/ml bleomycin for 3 h. Scale bars: 2 µm. D. Quantification of RAD51 foci on squashed protonema cells of wild type and BRCA2 complemented plants. Each dot presents the percentage RAD51-foci positive nuclei among DAPI stained cells from an image with the total cells counted in: wild type = 708, PpBRCA2* = 755, CpBRCA2 = 839 and AtBRCA2^ΔDBD^ = 709. Horizontal bars indicate the mean and standard deviation. The p-values shown were calculated using one-way ANOVA with Dunnett’s test for multi-comparison, ns; non-significant.

### 7. PpBRCA2 interacts with RAD51 through its BRC repeats

We next generated a *P. patens* line expressing GFP fused at the N-terminus of PpBRCA2 by inserting the *eGFP* sequence upstream of the *BRCA2* start codon (Fig. S1B). We determined the functionality of GFP-BRCA2 by analyzing plant growth, spore viability, meiosis, sensibility to bleomycin, HDR-dependent GT, and RAD51 focus formation. GFP-BRCA2 plants showed WT-like growth (Fig. S7A&B) and meiosis, with well-shaped spores (Fig. S7A) and a WT distribution of chromosome segregation (Fig. S7D). However, the spore viability was slightly reduced, suggesting a minor functional defect (Fig. S7C). Live imaging of germinated GFP-BRCA2 spores showed the GFP signal into the nucleus of the cell but did not mark all cells (Fig. 6A). This suggests that PpBRCA2 is detected as nuclear protein in cells, which may be undergoing a cellular division given its role in protecting stalled replication forks. Importantly, GFP-BRCA2 plants exhibited no hypersensitivity to bleomycin (Fig. S7E&F) and maintained an HDR-dependent GT efficiency at 14.4 %, close to the WT levels, with 98 % of cassette integration occurring at the targeted locus (Fig. S7I&J). Only a slight increase in RAD51-positive cells was observed after bleomycin treatment (Fig. S7G). Altogether, our analysis shows that GFP-BRCA2 is functional for DNA repair and HDR.

**Figure 6.**
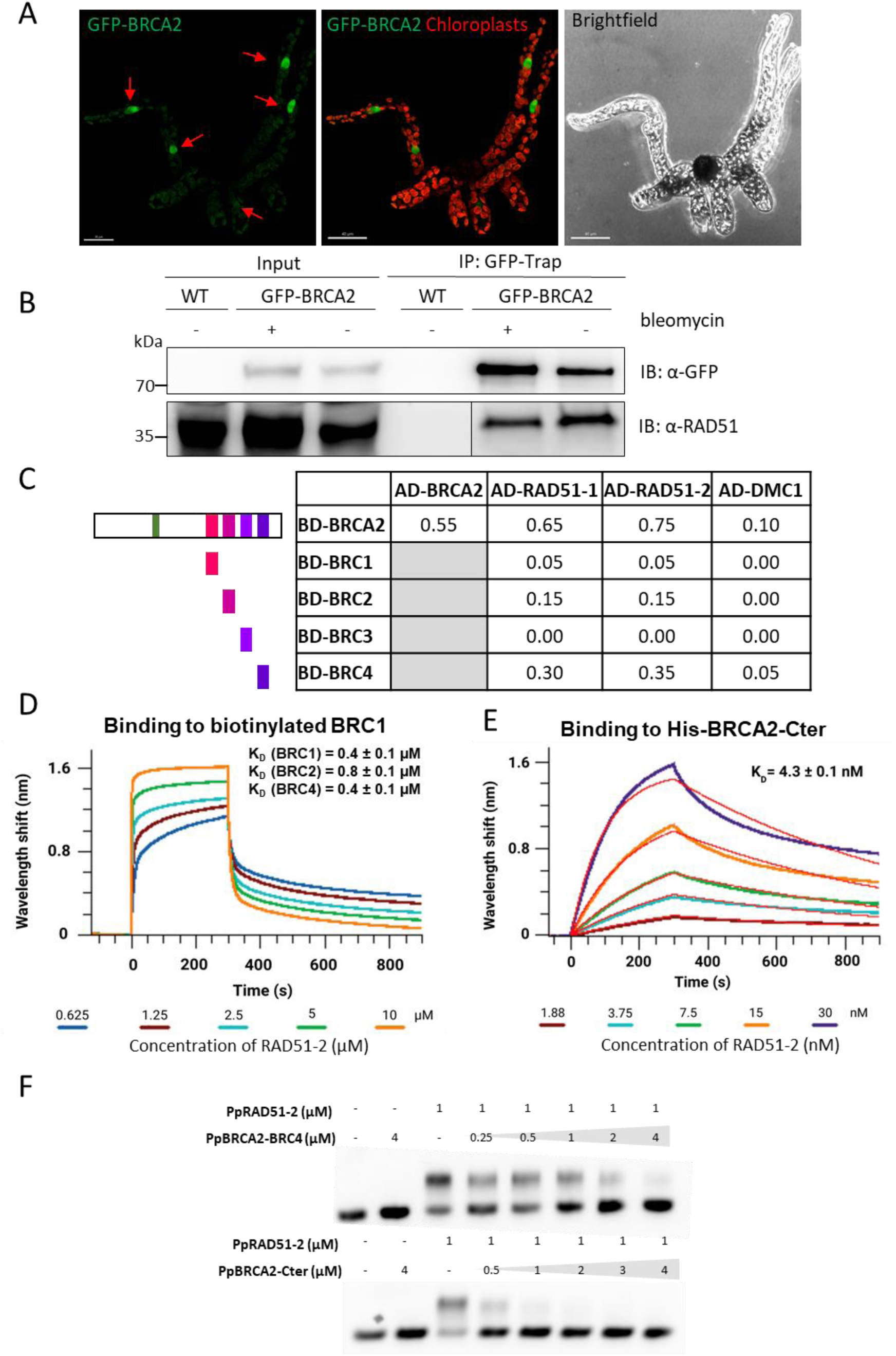
Cellular localization and interactions of PpBRCA2 protein. A. Representative image of a one-week-old germinated GFP-BRCA2 spore. Red arrows indicate nuclei expressing the strong GFP-BRCA2 signal in the nucleus. Chloroplasts’ autofluorescence is detected in red and light green. Scale bar: 30 µm. B. Co-immunoprecipitation of GFP-BRCA2 and PpRAD51. Total protein extracts of protonema from wild-type or GFP-BRCA2 line treated with or without bleomycin were immunoprecipitated with GFP-trap beads. Input and immunoprecipitated fractions were detected with anti-GFP and anti-RAD51 antibodies by Western blotting. C. The interaction between full-length PpBRCA2 or four moss BRC repeats and RAD51-1, RAD51-2, and DMC1 was evaluated using yeast two-hybrid assays. Each diploid was subjected to five-fold serial dilutions and spotted on SD/-LWH and SD/-LWHA media, where growing spots were assigned scores of 0.05 and 0.1, respectively. The final score for each diploid was obtained by summing the individual scores. A score of 0.75 shows the strongest interaction. Grey boxes indicate interactions that were not tested. BD; DNA binding domain, AD; activating domain. D. BioLayer Interferometry (BLI) interaction assays between biotinylated PpBRCA2-BRC peptides and PpRAD51-2. The BLI curves show PpBRCA2-BRC1 binding kinetics at increasing concentrations of PpRAD51-2 (0.625 to 10 µM). The Kd as a function of the PpRAD51-2 concentration was calculated by fitting the plateau shift of the association curve (steady-state mode; See Fig. S10A). Experiments with PpBRCA2-BRC2, -BRC3 and -BRC4 are displayed in Fig. S10C and replicates in Fig. S11A. No binding was observed with PpBRCA2-BRC3 (Fig S10B & S11A). E. BLI interaction assays between 6His-PpBRCA2-Cter and PpRAD51-2. The BLI curves show PpBRCA2-Cter binding kinetics at increasing concentrations of PpRAD51-2 (1.88 to 30 nM). Replicates are presented in Fig. S11B. F. EMSA assays indicate an inhibitory effect of PpBRCA2-Cter or PpBRCA2-BRC4 on RAD51 filament formation. The 3µM ssDNA (100 nucleotides) was incubated with 1 µM RAD51-2 and increasing concentrations (0 – 4 µM) of PpBRCA2-Cter or PpBRCA2-BRC4 for 15min at 37 °C followed by migration of agarose gel.

We then assessed whether the expression of GFP-BRCA2 was induced in response to DSB formation, similar to RAD51. Western blot analysis revealed no detectable change in the GFP-BRCA2 protein steady-state levels (Fig. S7H), suggesting that, unlike PpRAD51, PpBRCA2 is not upregulated following DSB formation. Given our AlphaFold3 analyses predicted high confidence interaction scores for BRC repeats (except BRC3) with PpRAD51, we tested these interactions using various approaches. Co-immunoprecipitation using GFP-trap beads confirmed a strong interaction between GFP-BRCA2 and RAD51 in moss independent of DSB formation (Fig. 6B). We also performed yeast two-hybrid (Y2H) assays to distinguish between the two *P. patens* RAD51 homologs (PpRAD51-1 and PpRAD51-2). Y2H assays detected a strong self-interaction of PpBRCA2 (Fig. 6C, Fig. S9), consistent with BRCA2 oligomerization observed in humans (79, 80). Additionally, the full-length PpBRCA2 strongly interacted with both PpRAD51 homologs (Fig. 6C, Fig. S9). AlphaFold3 predicted that the PpBRCA2 BRC repeats do not interact with PpDMC1 (Fig. S8A). Y2H assays consistently showed only weak interaction with PpDMC1 (Fig. 6C, Fig. S9). We then tested each BRC repeat individually against PpRAD51-1, PpRAD51-2, and PpDMC1 in Y2H assays. Overall, isolated BRC repeats showed weaker interactions compared to the full-length PpBRCA2 (Fig. S9). Among all BRCs, BRC4 displayed the strongest binding to both PpRAD51 proteins but a weak binding to PpDMC1 (Fig. 6C, Fig. S9). BRC1 and BRC2 exhibited moderate interactions with PpRAD51 but no detectable interaction with PpDMC1, while BRC3 failed to interact with both recombinases (Fig. 6C, Fig. S9), confirming AF predictions (Fig. 1C).

We next calculated the binding affinities to validate Y2H results using BioLayer Interferometry (BLI) assays with purified PpRAD51-2, PpDMC1 proteins, and BRC peptides. Biotinylated BRC peptides immobilized on a sensor were incubated with increasing concentrations of recombinant PpRAD51-2 or PpDMC1, followed by buffer immersion to monitor the association and dissociation kinetics (Kd). We measured Kd values of 0.4 µM, 0.8 µM, and 0.4 µM for PpRAD51-2 interactions with BRC1, BRC2 and BRC4, respectively (Fig. 6D, Fig. S10A&C, Fig. S11A). No detectable interaction was recorded between BRC3 and PpRAD51-2 (Fig. S10B, Fig. S11A) or between BRC repeats and PpDMC1 (Fig. S8B). Sequence alignments revealed a glycine-to-alanine substitution after the conserved FxxA motif in PpBRCA2-BRC3, potentially explaining its lack of interaction with PpRAD51. Replacing alanine with glycine restored PpRAD51 binding (Fig. S10D), confirming the importance of glycine residue. We then assessed the collective contribution of the BRC repeats with purified recombinant PpBRCA2-Cter, which contains all four BRCs at its C-terminal region. BLI assays exhibited a 100-fold stronger affinity of PpBRCA2-Cter for PpRAD51-2 (Kd = 4.3 nM) compared to individual BRC repeats (Fig. 6E & Fig. S11A), highlighting a cooperative binding effect among BRCs.

Structural studies of the human BRCA2 BRC4-RAD51 complex suggest that BRCA2 recruits monomeric RAD51 at DSB sites via its FxxA motifs (38). Excess human BRC4 peptides can disassemble preformed RAD51 filaments on ssDNA (81, 82). However, a ternary complex of truncated human BRCA2 containing all eight BRC repeats with RAD51, and ssDNA filaments is observed in the presence of non-hydrolyzable AMP-PNP (83). We determined the effects of PpBRCA2-BRC4 and PpBRCA2-Cter on preformed PpRAD51-ssDNA filaments by electrophoretic mobility shift assays (EMSA) to unravel how PpBRCA2 regulates RAD51-ssDNA filament dynamics. EMSA analysis showed that both PpBRCA2-Cter and BRC4 disrupt PpRAD51-ssDNA filaments in a concentration-dependent manner (Fig. 6F). However, complete filament disruption required different peptide concentrations: 2 µM for PpBRCA2-Cter and >4 µM for PpBRCA2-BRC4, confirming a significantly higher affinity of PpBRCA2-Cter for PpRAD51-2. These results highlight the cooperative action of BRC repeats in regulating RAD51 filament dynamics.

Taken together, PpBRCA2 plays a crucial role in HDR by mediating RAD51 recruitment through its BRC repeats. While individual BRCs show varying affinities, their combined action significantly enhances RAD51 binding, underscoring a conserved mechanism of BRCA2-mediated recombinase regulation. In contrast, PpBRCA2 exhibits minimal interaction with PpDMC1, suggesting a limited role in meiosis-specific recombination.

### 8. PpBRCA2 binds to ssDNA via its N-terminal region

Human BRCA2 binds to ssDNA through its canonical DBD as well as via its disordered N- and C-terminal regions (38, 41, 42). We explored if the disordered PpBRCA2 exhibits similar binding with various DNA substrates. We used purified recombinant PpBRCA2, PpBRCA2-Nter, and PpBRCA2-Cter to perform EMSA assays. Our EMSA analysis revealed that full-length PpBRCA2 binds to a 100 nt ssDNA and 150 nt-100 bp ss-dsDNA overhang but does not associate with 250 bp dsDNA (Fig. 7A). Further analysis identified that the N-terminal region of PpBRCA2 mediates ssDNA binding (Fig. 7B). To quantify binding affinities, we performed BLI analyses using biotinylated ssDNA immobilized on a sensor with increasing concentrations of recombinant PpBRCA2, PpBRCA2-Nter and PpBRCA2-Cter. PpBRCA2 and PpBRCA2-Nter strong ssDNA binding with dissociation constants (Kd) of 10-20 nM (Fig. 7C, Fig. S11B). In contrast, PpBRCA2-Cter displayed no detectable binding to ssDNA, consistent with the EMSA results (Fig. 7C, Fig. S11C). These findings demonstrate that the non-canonical PpBRCA2 binds to ssDNA through its N-terminal region.

**Figure 7.**
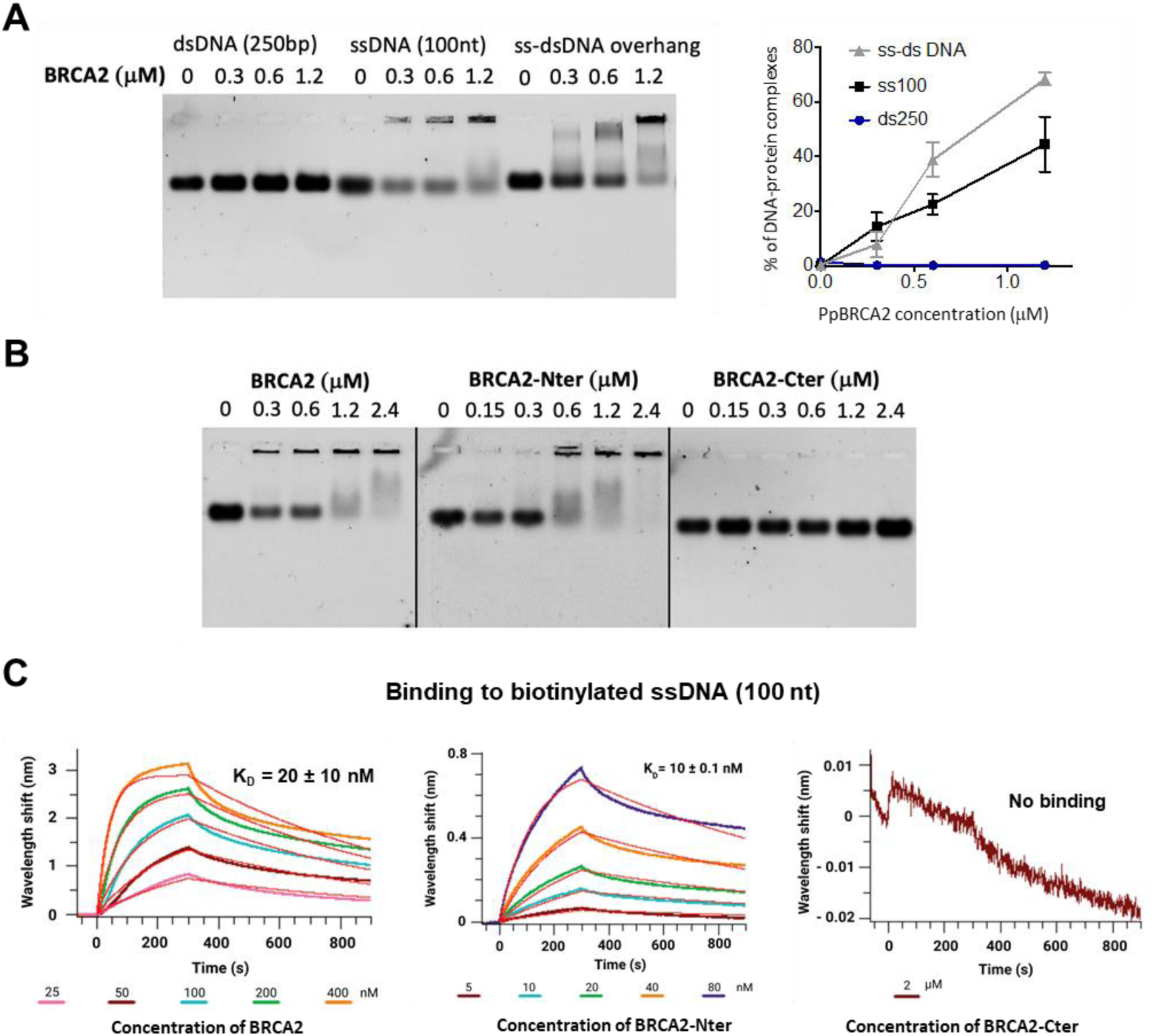
Interaction between PpBRCA2 and DNA by EMSA and BLI. A. Electrophoretic Mobility Shift Assay (EMSA) was performed by incubating 2 µM Cy5-labeled dsDNA (250 bp), ssDNA (100 nucleotides, nt) or ss-dsDNA (150 nt-100 bp) overhang substrates with increasing PpBRCA2 concentration (0 to 2.4 µM) for 7 min at 37°C followed by migration on agarose gel. The quantification of DNA-protein complexes showed that PpBRCA2 efficiently binds to ssDNA and ss-dsDNA overhang, but not to dsDNA. B. EMSA showed interaction of PpBRCA2 with ssDNA is mediated via its N-terminal. The increasing concentration of PpBRCA2, BRCA2-Nter or BRCA2-Cter (0 to 2.4 µM) was incubated with 2 µM ssDNA (100 nt) for 7 min at 37°C followed by migration on agarose gel. The PpBRCA2 and PpBRCA2-N-ter binds to ssDNA, but no binding was observed with PpBRCA2-Cter. C. BioLayer Interferometry (BLI) assays measuring the interaction between biotinylated ssDNA (100 nt) and PpBRCA2, PpBRCA2-Nter and PpBRCA2-Cter. The Kd values as a function of protein concentrations are obtained by fitting both the association and dissociation curves with replicates presented in Fig. S11B.

## DISCUSSION

We identified and characterized a unique plant BRCA2 homolog from *P. patens* (PpBRCA2), which has conserved BRC repeats but lacks the canonical DBD. Our study supports the existence of two types of BRCA2 homologs (Fig. 1E): (1) a canonical BRCA2, which includes a well-conserved folded DBD in most organisms, and (2) a non-canonical BRCA2, which lacks the canonical DBD as found in drosophila(3, 5) and mosses. *P. patens brca2Δ* mutants share similarities with canonical *brca2* plant mutants but also exhibit distinct phenotypes. While BRCA2 loss in mammals is lethal (10), *brca2Δ* mutants are viable in plants(4, 9) and drosophila(3, 5). *P. patens brca2Δ* mutants exhibit slow growth, with a ∼30% reduction in plant size, in contrast to the apparent normal growth observed in *A. thaliana brca2* (9). We provide compelling evidence that PpBRCA2 promotes DSB repair in somatic and meiotic cells. PpBRCA2 plays a critical role in DNA repair and genome stability by preventing hypersensitivity against genotoxic agents and limiting spontaneous mutation occurring in the genome, consistent with *P. patens rad51-1-2* phenotypes (71) and the elevated mutation rate observed in cells isolated from canonical *BRCA2-*mutated patients in humans (84). The non-canonical PpBRCA2 acts in somatic HDR as *brca2Δ* mutants exhibit a 10-fold reduction in GT, accompanied by an increase in random integrations. Further, the absence of RAD51 focus formation at DSBs in *P. patens brca2Δ* supports the role of PpBRCA2 as a RAD51 mediator. Loss of PpBRCA2 may either prevent RAD51 recruitment at DSB sites or lead to active RAD51 filament dissociation by anti-recombinase proteins such as FIGL1 (72). This suggests that PpBRCA2 is essential for both the nucleation and stabilization of RAD51 filaments.

Both *A. thaliana* and rice *brca2* mutants produce non-viable pollens and meiocytes fail to form bivalents owning to unrepaired breaks, leading to chromosome fragmentation detected from metaphase I onwards stages (4, 9). Similarly, *P. patens brca2Δ* mutants produce mostly non-viable spores and exhibit severe meiotic repair defects. Surprisingly, milder meiotic defects were observed at metaphase I in *P. patens brca2Δ* meiocytes. Only a small fraction of *P. patens brca2Δ* metaphase I cells displayed the bivalent shortage with a mix of univalent and bivalents, the remaining showed wild-type levels of bivalents. The *P. patens rad51-1-*2 meiocytes also displayed similar milder defects in bivalent formation with most cells having wild-type levels. However, a strong meiotic repair defect in terms of chromosome bridges and fragmentation was detected at anaphase I and subsequent stages. This indicates that BRCA2 and RAD51 are not essential for bivalent formation in moss, unlike their counterparts in Arabidopsis and rice. One speculation is that PpDMC1 can promote bivalents independent of BRCA2 and RAD51, though further work is necessary to confirm this hypothesis. Nonetheless, non-viable spores and chromosome breakage detected after metaphase I suggest critical roles of PpBRCA2 in RAD51-dependent meiotic DSB repair. Overall, the non-canonical PpBRCA2 mediates both somatic and meiotic RAD51 functions.

Canonical BRCA2 mediates HDR through its direct physical interaction with both RAD51 and DMC1 recombinases. We found a robust interaction between full-length PpBRCA2 and PpRAD51 homologs, with the BRC repeats binding to PpRAD51 at varying affinities. Isolated BRC1, 2, and 4 repeats showed micromolar affinities for RAD51, while BRC3 did not bind due to a glycine-to-alanine mutation following the FxxA motif. These findings are in concordance with previous studies showing varying interaction affinities of BRC repeats (34, 35). PpBRCA2-Cter (with all BRC repeats) displayed a 100-fold increase in binding affinity, reaching the nanomolar range, indicating a strong synergistic effect of the four BRCs on RAD51 binding. Our BLI data combined with the AF3 modeling suggest that PpBRCA2 BRC1, BRC2, and BRC4 repeats bind to monomeric RAD51-2, similar to human BRC4 repeat (38). Furthermore, the disruption of performed RAD51-ssDNA filaments by PpBRCA2-Cter and BRC4 suggests an inhibitory effect as previously observed with human BRC repeats (81, 82). Altogether, our data indicate that canonical and non-canonical BRCA2 orthologs regulate RAD51 loading and unloading through similar mechanisms.

Unlike canonical BRCA2 that strongly interacts with DMC1 (36, 85), PpBRCA2 BRC repeats bind to PpDMC1 with much lower affinity than to PpRAD51, supported by the low interaction scores from AF modeling. Despite a residual interaction in Y2H assays, no interaction with PpDMC1 was detected using in vitro assays. We propose that PpBRCA2 may have a conserved mediator function through RAD51 interaction in both somatic and meiotic cells and a less important role in DMC1-mediated recombination.

BRCA2 orthologs exist in eukaryotes, but their evolutionary dynamics across species remain puzzling, partly due to high sequence variability outside the BRC repeats and canonical DBD (86). In particular, the origin of non-canonical BRCA2 homologs during evolution is unresolved. One simplistic hypothesis is that they arose through the loss of exons encoding the DBD in certain lineages. Our analysis shows that canonical DBD loss is common to three moss genera – *P. patens*, *C. purpureus*, and *F. hygrometrica*. Although a lack of annotated genomes limits a broader analysis of DBD loss in other mosses, this finding suggests that the DBD loss likely occurred in the moss common ancestor after diverging from liverworts and hornworts. We propose that moss-specific evolutionary events led to the emergence of non-canonical BRCA2 within the bryophytes. Moreover, interactions involving the DBD, such as with DSS1, may not be essential for non-canonical BRCA2 functions. While DSS1 is crucial for canonical BRCA2 function(30), it appears dispensable in drosophila, where no genetic or protein interactions between drosophila BRCA2 and DSS1 have been observed (3). BRCA2 and DSS1 also do not appear to co-evolve as DSS1 homologs with additional cellular functions are present in yeasts that lack BRCA2. The role of two DSS1 homologs in *P. patens* remains to be explored in the context of HDR in the future.

Canonical BRCA2 homologs contain additional DNA-binding regions aside from the well-conserved and folded DBD. Human BRCA2 contains two such auxiliary regions at its N-terminal (NTD) and C-terminal (CTRB) (33, 41, 42). While the canonical DBD provides specificity for ssDNA binding, the auxiliary domains can bind both ssDNA and dsDNA (41, 42, 44). The current model suggests that human BRCA2 recruits RAD51 by either directly binding ssDNA through its canonical DBD or sliding on dsDNA via the auxiliary domains (44). The high frequency of pathogenic BRCA2 mutations within the canonical DBD in cancer patients underscores the importance of ssDNA binding of BRCA2 *in vivo* (46, 86). Surprisingly, deleting canonical DBD in mammalian BRCA2 does not cause lethality or completely disrupt HDR and RAD51 recruitment at DSBs, suggesting the auxiliary domains may compensate (48–51). Additionally, the presence of non-canonical BRCA2 homologs in drosophila and mosses suggests that the canonical DBD may not be essential for HDR in eukaryotes. How non-canonical BRCA2 compensates for the lack of a DBD remains unclear. Two possible scenarios are: (1) non-canonical BRCA2 operates independently of DNA binding, or (2) a cryptic domain ensures DNA binding activity. We demonstrate that non-canonical PpBRCA2 has an N-terminal domain that specifically binds ssDNA, outside the conserved BRC repeats. Our NMR analysis indicates that PpBRCA2 is an intrinsically disordered protein and lacks the well-folded globular domain found in canonical BRCA2. Our findings suggest that the N-terminal DNA binding activity of PpBRCA2 compensates for the loss of canonical DBD, reinforcing that the DNA binding activity, rather than the presence of canonical DBD, is a conserved feature among BRCA2 orthologs throughout evolution.

## ACKNOWLEDGMENTS

We thank Magali Nicaise-Aumont and Magali Noiray at I2BC for their assistance with the BLI experiments as well as Anne Wijkhuisen and the SPI (CEA Saclay) for providing access to their BLI instrument, as well as the French Infranalytics platform in Gif-sur-Yvette for NMR the Bruker 950 MHz spectrometer. The AlphaFold3 calculations were performed in collaboration with the Bioinformatics platform at I2BC. We are grateful to Mathilde Grelon at IJPB for the critical reading of the manuscript and discussions. This research was supported by the CNRS, the CEA-Saclay, the French Infrastructure for Integrated Structural Biology [ANR-10-INSB-05-01], and by ANR [ANR-22-CE12–0040] to PD, FN, SZJ and RK. This work has benefited from the support of IJPB’s cytology platforms. The IJPB benefits from the support of LabEx Saclay Plant Sciences-SPS [ANR-10-LABX-0040-SPS].

## AUTHOR’S CONTRIBUTIONS

A.Ch., A.Cr., S.A., U.T., S.Q., S.M., P.D., and P-F.P. produced data. A.Ch., S.Q., P.D., P-F.P. S.Z-J, F.N., and R.K. analyzed the data. P.D., S.Z-J, F.N., and R.K. conceived and designed the experiments. A.Ch., S.Q., S.Z-J, F.N., and R.K. wrote the manuscript with input from all authors.

**DATASET_S1**. HHpred search analysis using Arabidopsis AtBRCA2A as a query on human, drosophila, and moss proteomes.

**DATASET_S2**. HHpred search analysis using PpBRCA2 (Pp6C10_10830) as a query.

**DATASET_S3**. List of all primers, SgRNA, vectors, and peptides, along with raw data from genotoxic and mutator assays.

