## Supplementary material for "Moss BRCA2 lacking canonical DNA binding domain promotes efficient homologous recombination and binds to DNA": Supp figure

**Title:**

### SUPPLEMENTARY FIGURES

Figure S1

A

```
                                  BRC1                                                        BRC2
ss_pred       CCCceeecCCCccccCCHHHHHHHHHHHhcccCcccccccccCCcchhcCCCccccCCceecCCCCeeeCCHHHHHHHHHHhQ
AtBRCA2   66  EIPMFRTGLGKSVVLKESSIAKAKSILAEKVTYSDLRNTNCSIPQMRQVDTAETLPMFRTASGKSVPLKESSIAKAMSIL   145
              +...|+-++||...-+..+-.+|..+|+.+.                   ...+..-|+.++||...-+..+-.+|..||
PpBRCA2  249  NDNLFQLASGKRFQASAASKARAAALFESEV-------------------EPAGDSLFKLASGKQFQASAAAKARAAAMF   309
ss_pred       ccchhhhcCCcccccCHHHHHHHHHHHhCCC-------------------cccccchHhHHhhcccccccHHHHHHHHHHH
                                         BRC3
ss_pred        ccCcccCCCCccccCCCCccCccceeCCCCeeecCHHHHHHHHHHHccccc---cccccccccccccccccCCcccc
AtBRCA2   146  GSDKIIDSDNVLPRESGFGVSNSLFQTASNKKVNVSSAGLARAKALLGLEED---DLNGFNHVNQSSSSSQQHGWSGLKT   222
               +            ...-...-+.|..++||...-|.++-.||..||+.+.+   +..
PpBRCA2   310  E-------------SDVEPAGENLFNSASAKLFQASPAAKARAAALFESDVEPAGESL----------------------   354
ss_pred        h-------------ccCCcchHhHhhhhcccccccCHHHHHHHHHHHhcCCchHhHhH----------------------
                                                                         BRC4
ss_pred        cccccchhHhcccCCCccccccccCCccccCCCCCCCCCCceeeCCCCcCccCHHHHHHHHHHHhc
AtBRCA2   223  HEEFDATVVKHHSGTPGQYEDYVSGKRSEVLNPSLKVPPTKFQTAGGKSLSVSAEALKRARNLLG   287
                                                        |..|+||....|..|-.||..||+
PpBRCA2   355  -----------------------------------------FKLASGKPFQASAAAKARAAALFE   378
ss_pred        -----------------------------------------HHHhcCCccccCCHHHHHHHHHHHh
```

B

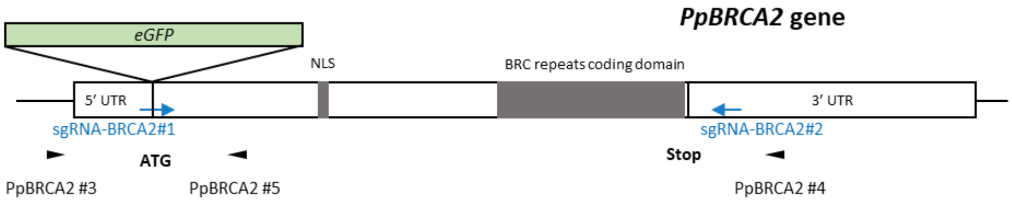


C

**> *Physcomitrium Patens*|Pp6C10_10830**

```
MMSMPTGFSVEQALEGGGFFPAQPVKISAHPSFPVAVDPVYVAPARPYPVIVNPFKPRVALREYVRRLSGENKENDIGKEQMDSPSGLPKTKT
LGLRRRHSGPPSSVVLGAQSEGVARDKRKFNAPRQTPVKRQREETNEDVNGDACQSLPFASVKPMEVEQSLGSSPSNRQPAESLTPSFKRHSQ
ESFSPINRRPLAPLNFSTNRQVTELHTPITKPQSTERFMPPIHQIPSKPQFSGGPECGGSALNDNLFQLASGKRFQASAASKARAAALFESEV
EPAGDSLFKLASGKQFQASAAAKARAAAMFESDVEPAGENLFNSASAKLFQASPAAKARAAALFESDVEPAGESLFKLASGKPFQASAAAKAR
AAALFESDADAMDHMSFDD
```

**>*Funaria hygrometrica*|Fh_23015**

```
MMSMPTGFSVELALEGGVFIPAQPVKNSAHCSFPVAVDPVHVAPARPYPVLANPFKPRFALREYVRRLSGENKENDIAKEQMDSPSGLPKTKT
LGLRRRHSGPPSSVVLGAQNEGVARDKRKFNAPRQTPVKRQREETNEDMNGDVCQALPYASTKSMEVEQTLGTSPSNQQPAESLTPSFKRHLH
EPFSPINRRPLAPLNFSTNRQVVEPHTPTTKPFTERFMSPIHQLPSKPQFSGGLGSGGSGPNDNLFQSASGKRFQASAASKARAAVLFESEDE
PAGDSLFQLASGKQFQASAAAKARAAAMFESDVEPAGESLFKLASGKPFQASAAAKARAAALFESDADPMDHVSFDN
```

**>*Ceratodon purpureus*|CepurR40.9G141900.1.p**

```
MSGREVDAEGGGMRRGEDCVEVGVQASGVDRVAPCDGAGVRSPYPAVVDPFKRRVALREYVKQLRGDNKENERGKDSEVSEKSPVSGGVVKKT
LGLCRRHVGGVAAVAVSREGMLRDKRKFNAPRQTPEKRQRPDQVDVSPGTELSRMESRMVVEEGGEEMSGGMRQPVFSFASREHVQLELSPGQ
VSGGDGGDGVAGARWSPYGGRDSFLSPAPSPAPSPSRQAQLPESYTLPFKWQAPEPKTPTIRKPLAPVNIPVNRQTVDLHPSPMDQPRPNFSG
SLERNSSGSGGSLFQTASGKPFQASAAAKARAALLFESDAELADFFAMEL
```

D E

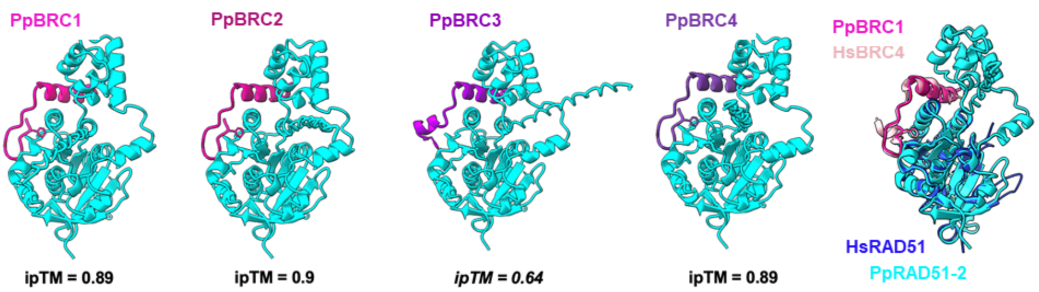


F

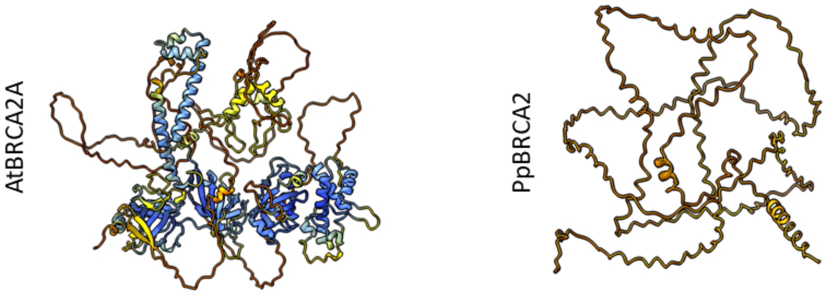

**Figure S1. Protein sequences and structural models of BRCA2 homologs**

A. The HHpred alignment of *Arabidopsis thaliana* BRCA2 (66 to 287 aa) and *Physcomitrium patens* BRCA2 (249 – 378 aa) showing conservation of four BRC repeats marked in in pink, orchid, violet and purple colors.

B. Schematic representation of the genomic locus of *Pp6C10_10830 (PpBRCA2* gene) with 5' and 3' UTRs, start and stop codons as well as NLS and BRC repeat region indicated in grey. Positions of *eGFP* insertion (inverted triangle) in green, guide RNAs in blue and genotyping primers in black are indicated.

C. Sequence of BRCA2 proteins from *Physcomitrium patens, Funaria hygrometrica* and *Ceratodon purpureus*. In *P. patens* BRCA2, NLS is highlighted in green and four BRC repeats are highlighted in pink, orchid, violet and purple colors. In *F. hygrometrica* and *C. purpureus*, BRC repeats are highlighted on orchid color.

D. Predicted 3D structure models of interaction between PpRAD51-2 (cyan) and four BRC repeats built with the AlphaFold3 program, with ipTM values > 0.7 representing high-confidence scores except for BRC3. The conserved phenylalanine of the FxxA motif from BRC1, 2 and 4 is buried in the hydrophobic pocket of RAD51, which is not the case for BRC3 interaction.

E. Overlay of the model of the complex between PpRAD51-2 (cyan) and the BRC1 repeat of PpBRCA2 (pink) built with the AlphaFold3 program and the crystal structure of the chimeric HsRAD51-HsBRC4 protein (PDB: 1n0w). HsRAD51 is displayed in blue and HsBRC4 in light pink. The conserved phenylalanine of the FxxA motif binding to RAD51 pocket is shown.

F. Alphafold3 structure models of a canonical BRCA2 (*A. thaliana* AtBRCA2) with a well folded DBD and a non-canonical BRCA2 (*P. patens* PpBRCA2) without a DBD.

Figure S2

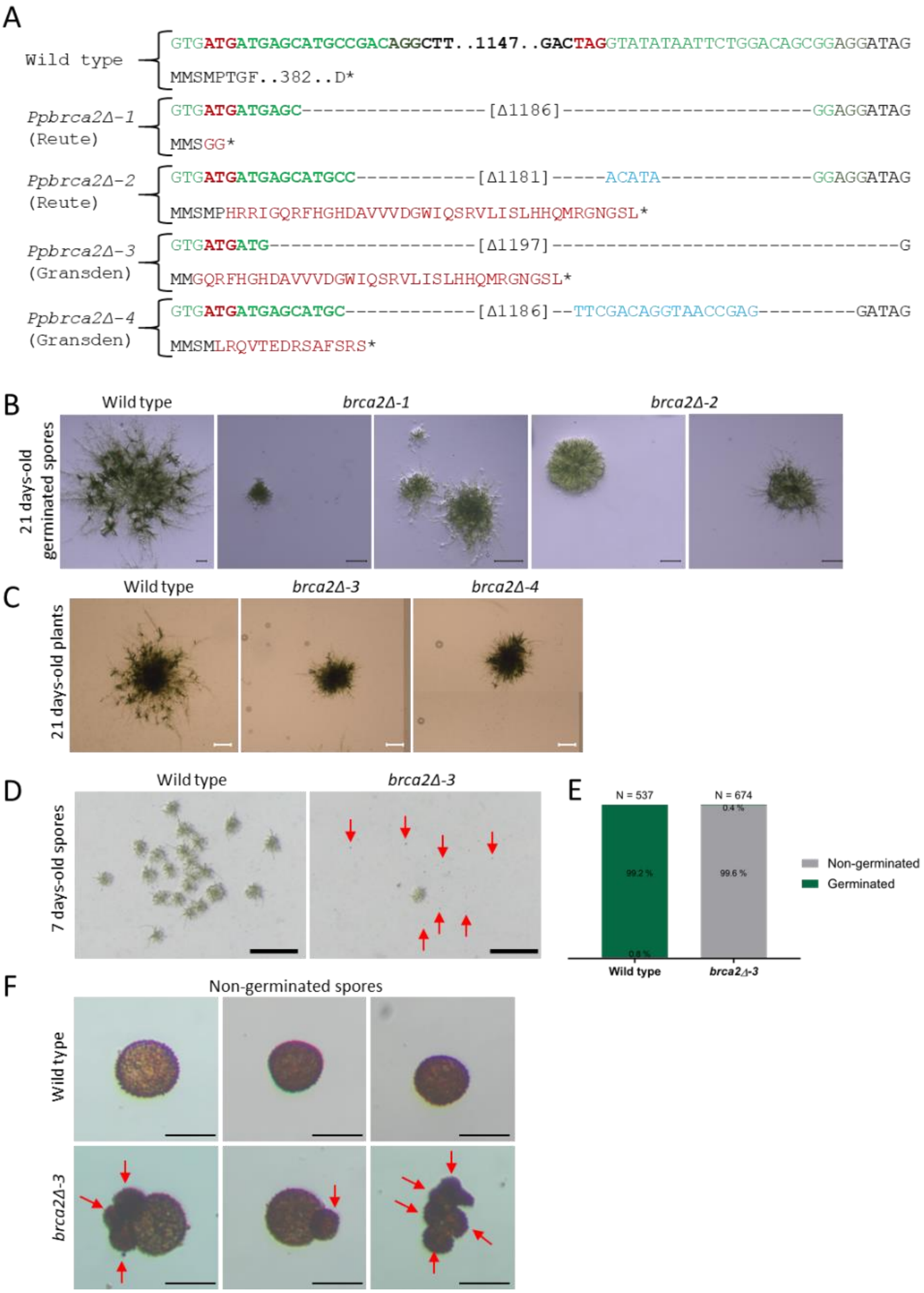

A
Wild type
GTGATGATGAGCATGCCGACAGGCTT..1147..GACTAGGTATATAATTCTGGACAGCGGAGGATAG
MMSMPTGF..382..D*
Ppbrca2Δ-1
(Reute)
GTGATGATGAGC-----------------[Δ1186]-----------------------GGAGGATAG
MMSGG*
Ppbrca2Δ-2
(Reute)
GTGATGATGAGCATGCC-----------[Δ1181]-----ACATA--------------GGAGGATAG
MMSMPHRRIGQRFHGHDAVVVDGWIQSRVLISLHHQMRGNGSL*
Ppbrca2Δ-3
(Gransden)
GTGATGATG-------------------[Δ1197]-------------------------------G
MMGQRFHGHDAVVVDGWIQSRVLISLHHQMRGNGSL*
Ppbrca2Δ-4
(Gransden)
GTGATGATGAGCATGC------------[Δ1186]--TTCGACAGGTAACCGAG---------GATAG
MMSMLRQVTEDRSAFSRS*
B
21 days-old germinated spores
Wild type
brca2Δ-1
brca2Δ-2
C
21 days-old plants
Wild type
brca2Δ-3
brca2Δ-4
D
7 days-old spores
Wild type
brca2Δ-3
E
N = 537
N = 674
99.2 %
0.8 %
99.6 %
0.4 %
Non-germinated
Germinated
Wild type
brca2Δ-3
F
Non-germinated spores
Wild type
brca2Δ-3

**Figure S2. Growth, spore germination, and viability defects of *P. patens brca2Δ* mutants**

A. DNA sequence of the *PpBRCA2* locus in wild-type and four *brca2*Δ mutants (*brca2*Δ*-1* and *brca2*Δ-*2* Reute ecotype, and *brca2*Δ*-3,* and *brca2*Δ*-4* in Gransden ecotype) is shown, with start and stop codons in red, and guide RNA sequences in green with PAM in bold. In the wild type, a dotted line with numbers indicates nucleotide present between guides. Dashed lines with bracketed numbers show deleted sequences in *brca2*Δ mutants, with insertions marked in blue. The amino acid sequences are presented in single-letter code: wild-type sequence in black, and altered sequence in red in *brca2*Δ.

B. Images of wild-type and *brca2Δ* mutant germinated spores after 21 days showed aberrant growth development in the absence of PpBRCA2. Scale bars: 1 mm.

C. Representative images of 21-day-old wild-type, *brca2Δ-3,* and *brca2Δ-4* mutant plants in the Gransden ecotype showed a reduction of plant diameter in the absence of PpBRCA2. Scale bars: 1mm.

D. Representative images of 6-days old wild-type and *brca2Δ* germinated spores in the Grandsen ecotype. A selection of non-germinated spores is indicated with red arrows. Scale bars: 1mm.

E. Quantification of spore germination rate after 6 days of growth. Spores are coming from one single capsule of wild type and *brca2Δ* in the Gransden ecotype.

F. Three images of wild-type and *brca2Δ* non-germinated spores in the Gransden ecotype. Red arrows indicate anomalies in the shape and size of mutant spores. Scale bars: 30 µm.

Figure S3

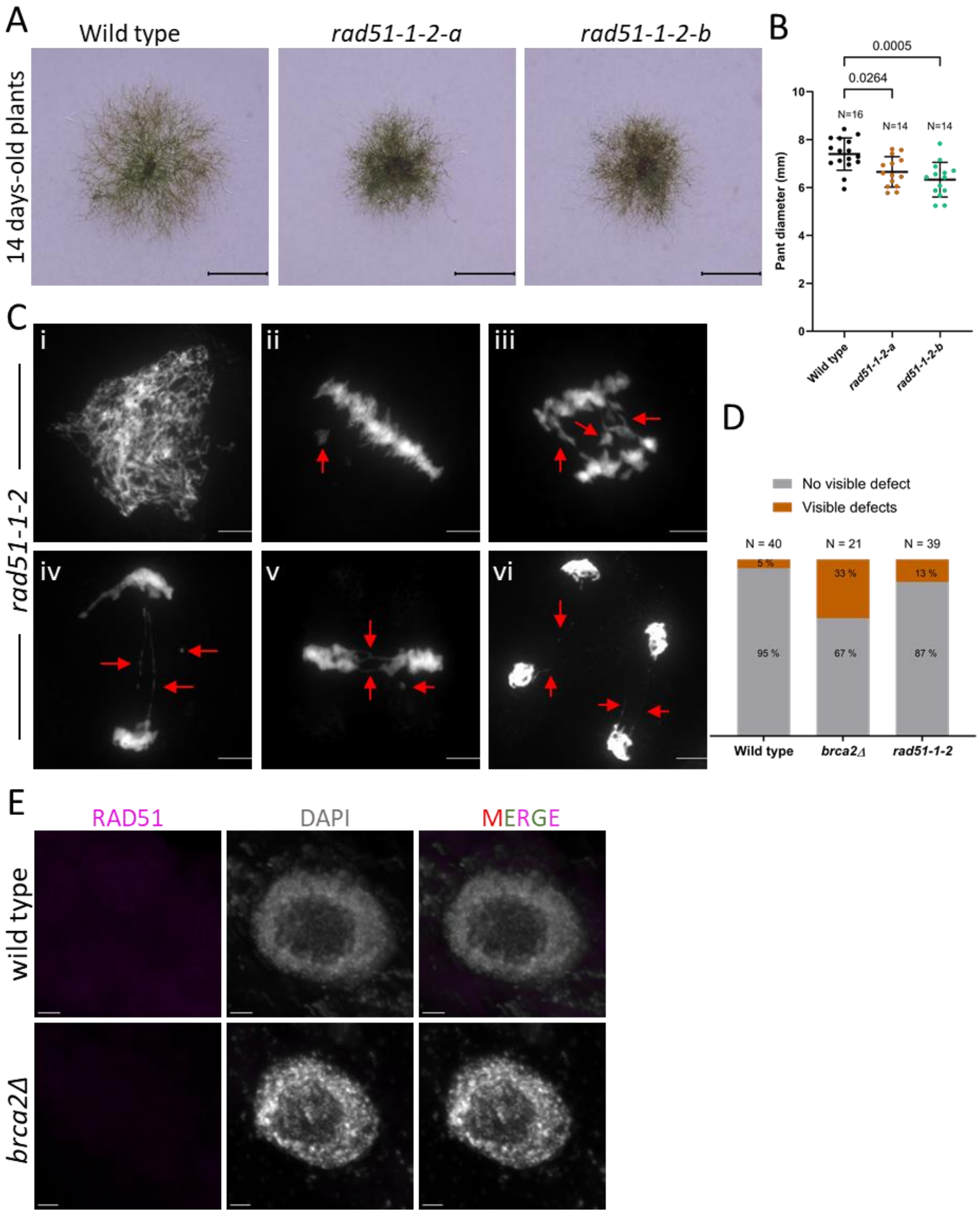


**Figure S3. Somatic and meiotic defects in *P. patens rad51-1-2* double mutants.**

A. Representative images of 14-day-old wild-type, *rad51-1-2-a* and *rad51-1-2-b* plants on minimal media. *rad51-1-2-a* and *rad51-1-2-b* are two different mutant alleles. Scale bars: 1 mm.
B. Quantification of the size of 14-day-old plants with each color dot representing an individual plant along with mean and standard deviation indicated by horizontal bars. Wild-type data is used from Fig.2. The p-values were computed using one-way ANOVA with Dunnett's test for multi-comparison.
C. Representative images of DAPI-stained chromosome spreads at leptotene (i), metaphase I (ii), anaphase I (iii), telophase I (iv), metaphase II (v) and telophase II (vi) in *rad51-1-2* meiocytes. Red arrows indicate lagging chromosomes, DNA fragmentation or univalent chromosomes. Scale bars: 5 µm.
D. Histogram comparing defects in metaphases I of wild type, *brca2*Δ, (Fig.2) and *rad51-1-2* mutant.
E. Immunolocalization of RAD51 (purple) in nuclei of the squashed protonema cells of wild type and *brca2Δ* plants without bleomycin treatment. Scale bars: 2 µm.

Figure S4

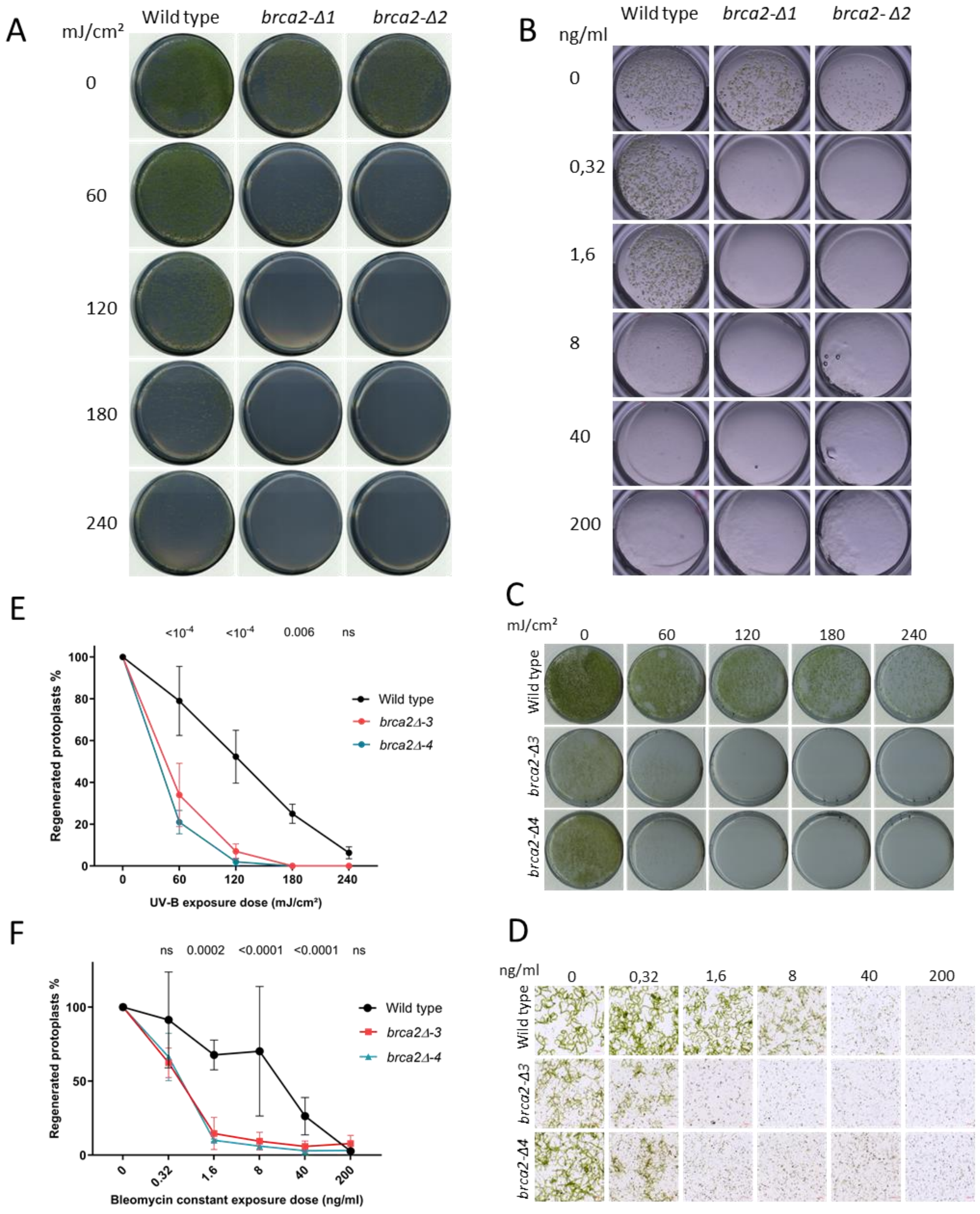
A
mJ/cm²
Wild type
brca2-Δ1
brca2-Δ2
0
60
120
180
240
B
ng/ml
Wild type
brca2-Δ1
brca2- Δ2
0
0,32
1,6
8
40
200
E
<10⁻⁴ <10⁻⁴ 0.006 ns
Regenerated protoplasts %
UV-B exposure dose (mJ/cm²)
Wild type
brca2Δ-3
brca2Δ-4
C
mJ/cm²
0 60 120 180 240
Wild type
brca2-Δ3
brca2-Δ4
F
ns 0.0002 <0.0001 <0.0001 ns
Regenerated protoplasts %
Bleomycin constant exposure dose (ng/ml)
0 0.32 1.6 8 40 200
Wild type
brca2Δ-3
brca2Δ-4
D
ng/ml
0 0,32 1,6 8 40 200
Wild type
brca2-Δ3
brca2-Δ4

**Figure S4. Hypersensitivity to genotoxic agents in wild-type and *brca2Δ* mutants**

A. Representative images of regenerated protoplasts from wild-type, *brca2Δ-1* and *brca2Δ-2* in the Reute ecotype at 21 days after exposure to UV-B.

B. Representative images of regenerated protoplasts from wild-type, *brca2Δ-1* and *brca2Δ-2* in the Reute ecotype at 21 days after exposure to bleomycin.

C. Representative images of regenerated protoplasts from wild type, *brca2Δ-3,* and *brca2Δ-4* in the Gransden ecotype at 21 days after exposure to UV-B.

D. Representative images of regenerated protoplasts from wild type, *brca2Δ-3,* and *brca2Δ-4* in the Gransden ecotype at 21 days after exposure to bleomycin.

E. Survival curves of wild-type, *brca2Δ-3* and *brca2Δ-4* protoplasts regenerating 6 days after exposure to UV-B. Values are normalized on the non-treated sample. For each point, the mean of three repeats is plotted with error bars indicating standard deviation. The p-values shown were calculated using two-way ANOVA with Dunnett's test for multi-comparison. ns; non-significant.

F. Survival curves of wild-type, *brca2Δ-3* and *brca2Δ-4* protoplasts regenerating 6 days after exposure to bleomycin. Values are normalized on the non-treated sample. For each point, the mean of three repeats is plotted with error bars indicating standard deviation. The p-values shown were calculated using two-way ANOVA with Dunnett's test for multi-comparison. ns: non-significant.

Figure S5

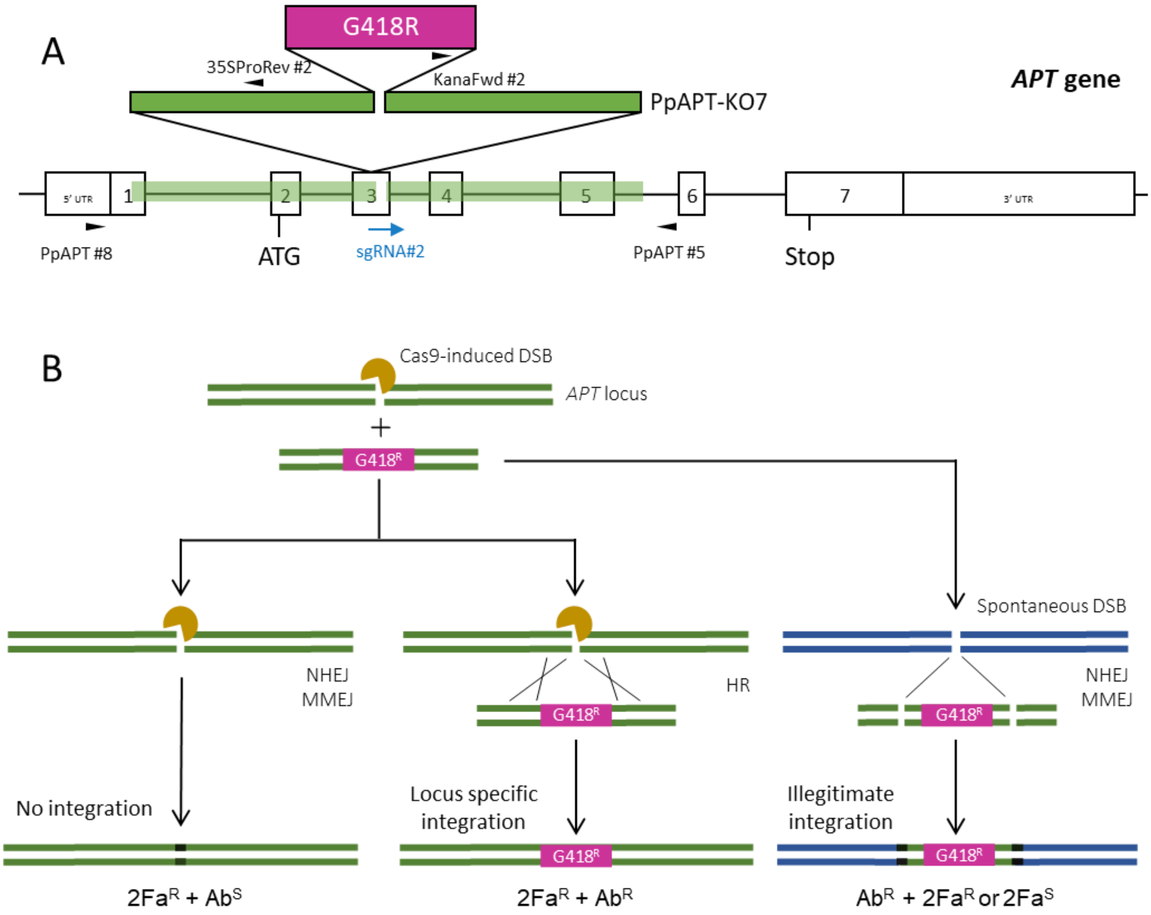


**Figure S5. Schematic representation of gene targeting strategy using the *APT* reporter gene**

A. Structure of *APT* gene with 7 exons and 5'&3' UTR in open boxes along with start and stop codon. DSB site is indicated by sgRNA#2 in blue and primer positions are shown by black arrowheads. Homology regions in the *APT* gene and the donor cassette (PpAPT-KO7) are highlighted in green. Inverted triangles indicate the insertion site.

B. Three scenarios are expected from the DSB repair with donor cassette (PpAPT-KO7). (1) Plants with no integration at the *APT* locus aroused from the Cas9-DSB repair by non-HDR pathways such as NHEJ, leading to 2FA resistance and G418-sensitivity. (2) Plants with the G418 cassette integration at the *APT* locus-specific resulted from homologous recombination repair, conferring resistance to 2FA and G418. (3) Plants with the G418 cassette illegitimate integration in the genome were resistant to G418, along with or without 2FA resistance resulting from inaccurate repair of breaks in the *APT* gene by non-HDR.

Figure S6

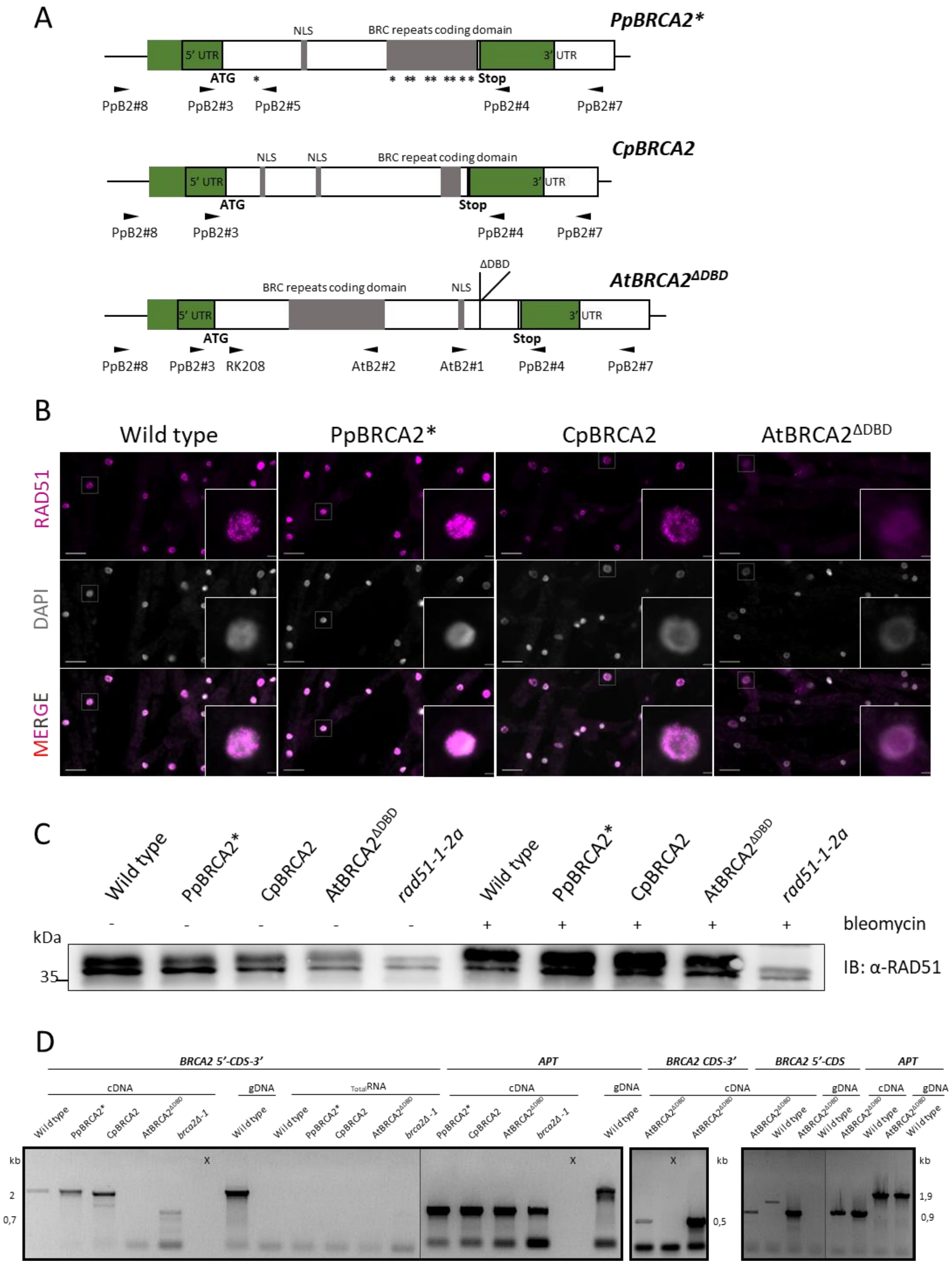
A
PpBRCA2*
NLS
BRC repeats coding domain
5' UTR
3' UTR
ATG
Stop
PpB2#8
PpB2#3
PpB2#5
PpB2#4
PpB2#7
CpBRCA2
NLS
NLS
BRC repeat coding domain
AtBRCA2ΔDBD
BRC repeats coding domain
NLS
ΔDBD
RK208
AtB2#2
AtB2#1
B
Wild type
PpBRCA2*
CpBRCA2
AtBRCA2ΔDBD
RAD51
DAPI
MERGE
C
rad51-1-2a
bleomycin
kDa
35
IB: α-RAD51
D
BRCA2 5'-CDS-3'
APT
BRCA2 CDS-3'
BRCA2 5'-CDS
cDNA
gDNA
TotalRNA
brca2Δ-1
kb
2
0,7
0,5
1,9
0,9

**Figure S6. Complementation of *P. patens brca2*Δ*-1* mutant**

A. Schematic representation of the *PpBRCA2*, CpBRCA2,* and *AtBRCA2*$^{\Delta DBD}$ alleles complementing the *PpBRCA2* locus in *P. patens*. Homology arms are highlighted in green with indicated positions of 5'& 3' UTR, start & stop codons, NLS & BRC repeat regions (in grey), and genotyping primers (black arrowheads). Silent substitutions in *PpBRCA2** are indicated with *.

B. RAD51 immunostaining in magenta and DAPI-stained nuclei from the wild-type, PpBRCA2*, CpBRCA2, and AtBRCA2$^{\Delta DBD}$ squashed protonema cells treated with 5 µg/ml bleomycin for 3 h. Scale bars: 20 µm. Single nuclei for each image are shown in the inset. Scale bars: 2 µm.

C. Western blot analysis of RAD51 steady-state-levels with anti-RAD51 antibody from the total protein extracts of wild-type, PpBRCA2*, CpBRCA2, and AtBRCA2$^{\Delta DBD}$ protonema treated with or without 5 µg/ml bleomycin for 3 hours.

D. RT-PCR expression analysis of wild-type, PpBRCA2*, CpBRCA2, and AtBRCA2$^{\Delta DBD}$ transcripts. PCRs were performed on cDNA prepared from total RNA extracted from respective genotypes. Genomic DNA and total RNA without RT were used as negative controls for RT reaction, while *APT* gene-specific PCRs served as positive controls. BRCA2 5'-CDS-3' denotes PCR amplifying the whole coding sequence. BRCA2 5'-CDS or BRCA2 CDS-3' indicates amplification of either 5' or 3' region of the *AtBRCA2* coding sequence.

Figure S7

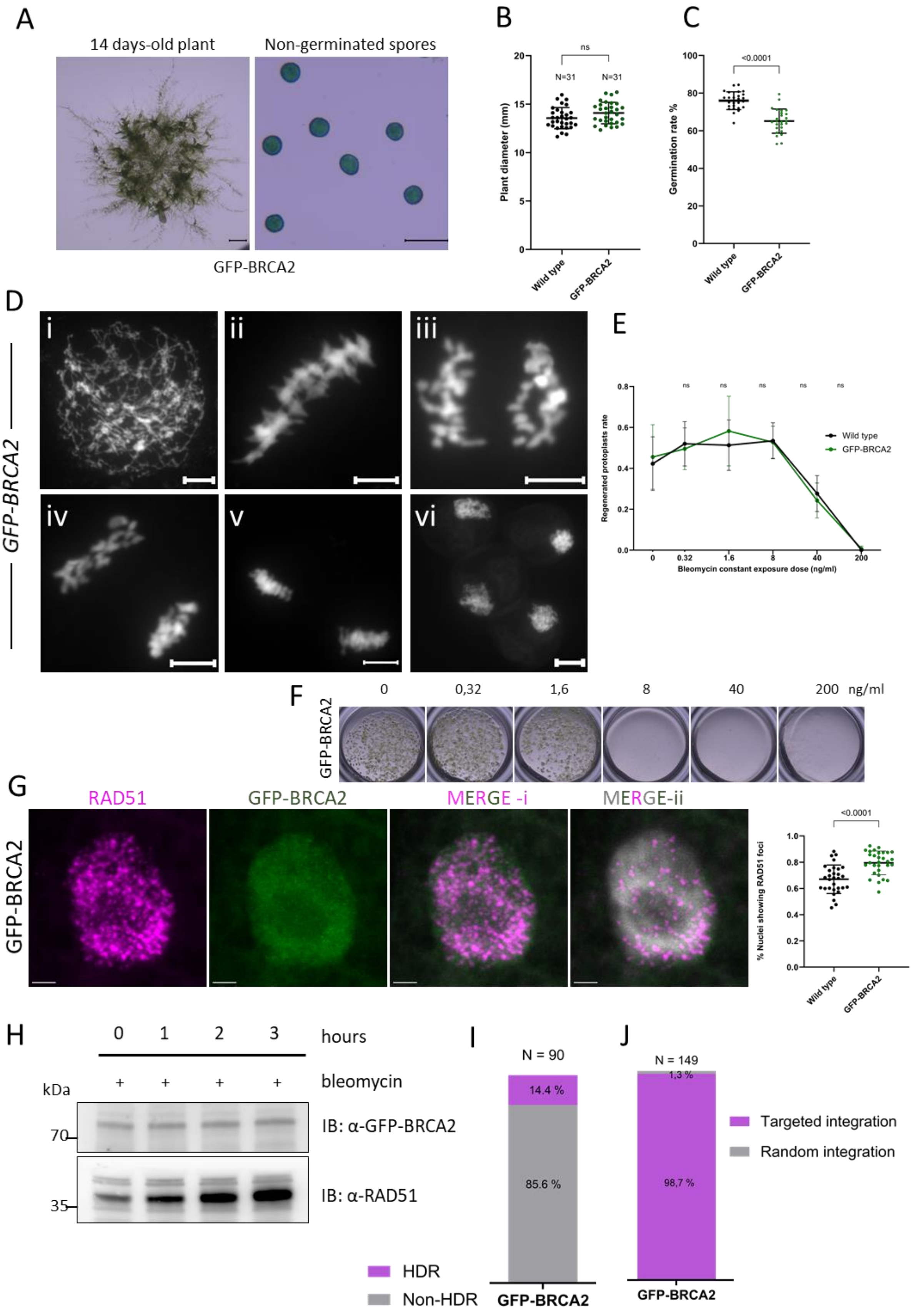

A
14 days-old plant
Non-germinated spores
GFP-BRCA2
B
ns
N=31
N=31
Plant diameter (mm)
Wild type
GFP-BRCA2
C
<0.0001
Germination rate %
Wild type
GFP-BRCA2
D
GFP-BRCA2
i
ii
iii
iv
v
vi
E
Regenerated protoplasts rate
Bleomycin constant exposure dose (ng/ml)
Wild type
GFP-BRCA2
F
GFP-BRCA2
0
0,32
1,6
8
40
200 ng/ml
G
GFP-BRCA2
RAD51
GFP-BRCA2
MERGE -i
MERGE-ii
<0.0001
% Nuclei showing RAD51 foci
Wild type
GFP-BRCA2
H
0 1 2 3 hours
+ + + + bleomycin
kDa
70
35
IB: α-GFP-BRCA2
IB: α-RAD51
I
N = 90
14.4 %
85.6 %
GFP-BRCA2
HDR
Non-HDR
J
N = 149
1,3 %
98,7 %
GFP-BRCA2
Targeted integration
Random integration

**Figure S7. Analysis of GFP-fusion functionality in the GFP-BRCA2 plants**

A. Representative image of a 14-day-old plant on minimal media and Alexander-stained non-germinated spores from the GFP-BRCA2 line. Scale bars: 1 mm. & 50 µm.
B. Quantification of 14-day-old plant diameter on minimal media, with each dot representing an individual plant and horizontal bars indicating the mean and standard deviation. The p-values calculated using parametric student test, ns; non-significant.
C. Quantification of percent spore germination after 6 days of growth. Spores were pooled from seven (GFP-BRCA2) or eight (wild type) different capsules, with each dot reporting the percentage of germination of a group of 8 to 25 spores adding up to the following total spore numbers: wild type = 5099 and GFP-BRCA2 = 2822. Horizontal bars indicate mean and standard deviation. The p-values shown were calculated using parametric student test.
D. Representative images of DAPI-stained chromosome spreads at leptotene (a), metaphase I (b), anaphase I (c), telophase I (d), metaphase II (e) and mature spores (f) in GFP-BRCA2 meiocytes. Scale bars: 5 µm.
E. Survival curves of regenerated protoplasts from wild-type (same data as in Fig.3) and GFP-BRCA2 after 3 days of constant exposure to bleomycin. Values are independent. Each point is the mean of three independent experiments plotted with error bars indicating standard deviation. The p-values shown were calculated using two-way ANOVA with Dunnett's test for multi-comparison, ns; non-significant.
F. Representative images of GFP-BRCA2 regenerated protoplasts after 14 days of constant exposure to bleomycin.
G. Immunolocalization of RAD51 (purple) and GFP (green) on the squashed protonema cells of GFP-BRCA2 plants treated with 5 µg/ml bleomycin for 3 h. Merged images of GFP-BRCA2 and RAD51 signals as well as with DAPI are shown. The percentage of RAD51-positive cells from wild-type (N= 1698) and GFP-BRCA2 (N=1232) squashed protonema after bleomycin treatment is presented in the scatter plot with the mean and standard deviation and the p-values computed using parametric student test. Scale bars: 2 µm.
H. Western blot analysis of GFP-BRCA2 and RAD51 steady-state-levels with anti-GFP and anti-RAD51 antibodies from the total protein extracts of GFP-BRCA2 protonema treated with or without 5 µg/ml bleomycin for 3 hours.
I. Percentage of gene targeting events (donor G418 cassette insertion) by HDR (purple) and non-HDR events (no insertion) in grey at the *APT* locus in *GFP-BRCA2* protoplasts. Cassette integrations were confirmed by genotyping $G418^R$ and $2\text{-}FA^R$ plants using *APT*-locus-specific primers.
J. Percentage of targeted (purple) and random integration (grey) of the donor G418 cassette at the *APT* locus in *GFP-BRCA2* protoplasts. Cassette integrations were confirmed by genotyping stable $G418^R$ plants using *APT*-locus-specific primers.

Figure S8

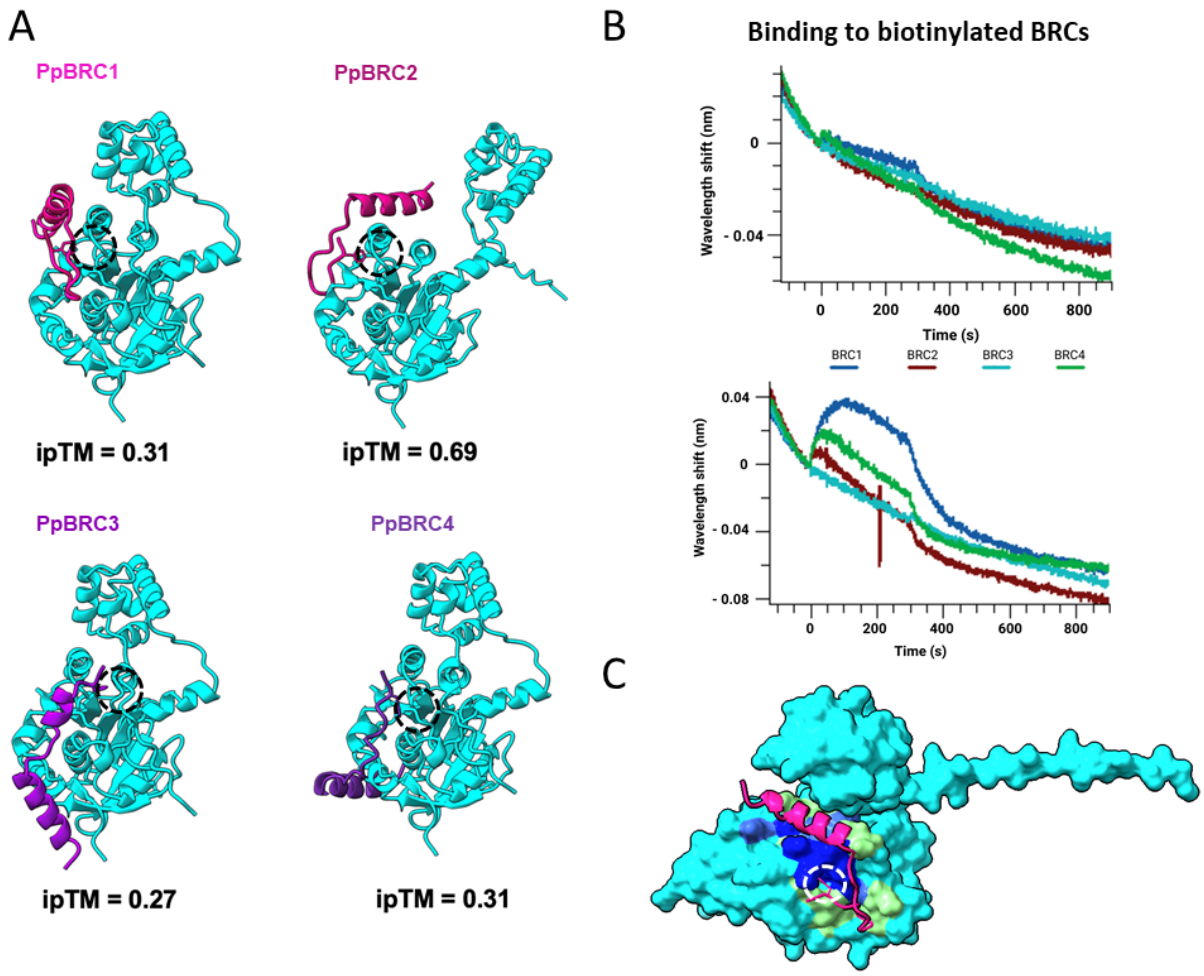


**Figure S8. Alphafold modeling and BioLayer Interferometry (BLI) interaction between BRC repeats and PpDMC1**

A. 3D structure models of interaction between PpDMC1 (cyan) and PpBRCA2-BRC repeats marked in in pink, orchid, violet and purple built with AlphaFold3. The conserved phenylalanine of the FxxA motif in BRC repeats is indicated by dotted circles with the ipTM scores shown below each model.
B. BLI assays between biotinylated PpBRCA2-BRC1, -BRC2, -BRC3, -BRC4, and PpDMC1 (10 μM), failed to detect any interaction.
C. 3D structure model of the interaction between PpDMC1 (cyan) and the PpBRCA2-BRC1 (pink) built with AlphaFold3. The residues of the model colored differently represent the PpRAD51-2 interaction site with BRC1 and display the conserved and non-conserved residues with PpDMC1 (light green: conserved residues, royal blue: conserved substitutions, and blue: non-conserved substitutions). The conserved phenylalanine of the FxxA motif in BRC repeats is indicated by a white dotted circle.

Figure S9

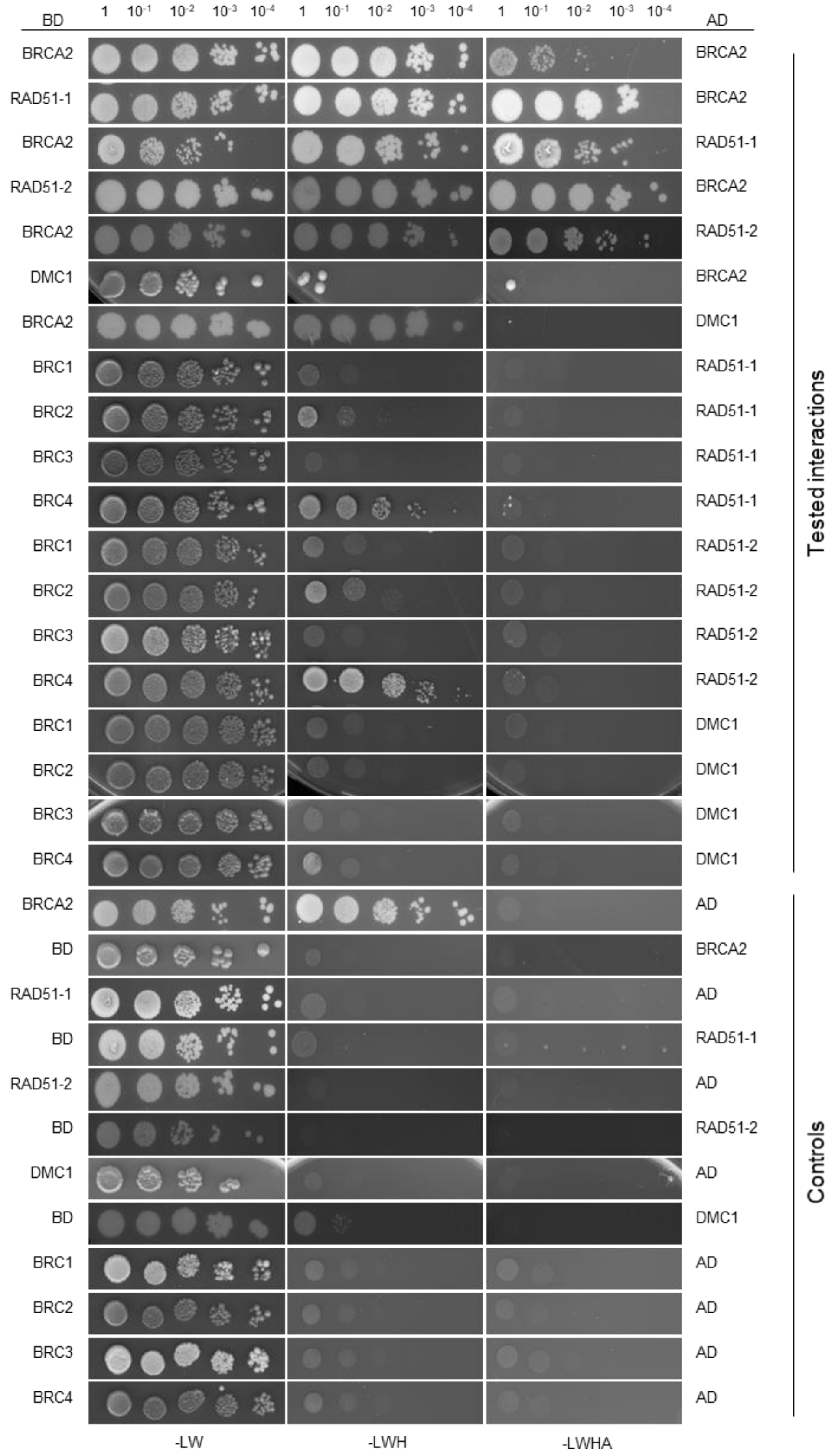


**Figure S9. Yeast-two hybrid interaction assays between PpBRCA2, PpRAD51 and PpDMC1**
Five-fold serial dilutions of diploid yeast strains harboring the indicated fusion proteins were spotted on –LW, -LWH, and –LWHA media. BD, DNA binding domain; AD, Activating domain.

Figure S10

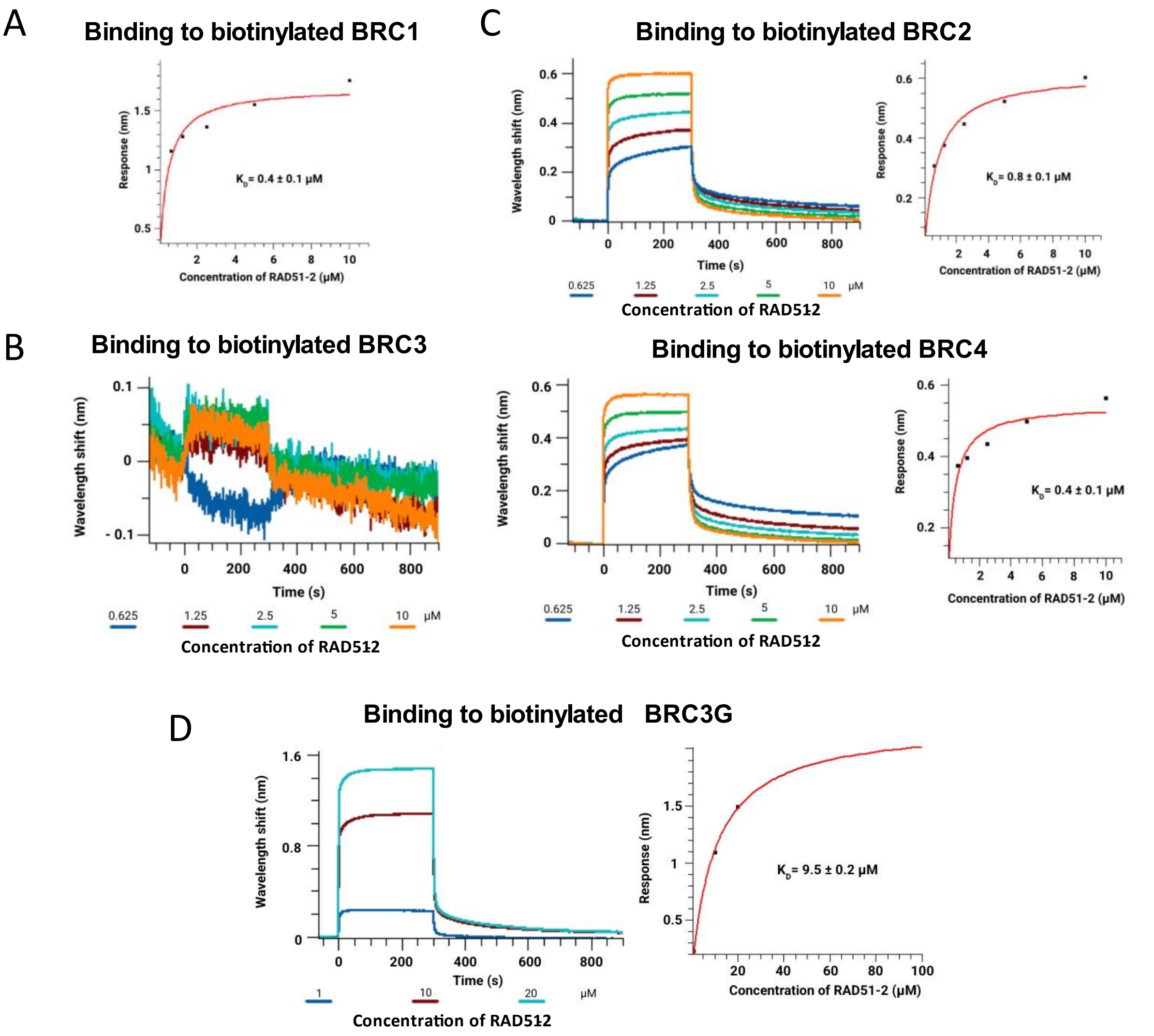


**Figure S10. BioLayer Interferometry (BLI) curves corresponding to the interaction between biotinylated PpBRCA2-BRC repeats and PpRAD51-2**

A. Interaction between PpBRCA2-BRC1 and PpRAD51-2. The Kd as a function of the PpRAD51-2 concentration is obtained by fitting the plateau shift of the association curve (steady-state mode).

B. Interaction between PpBRCA2-BRC3 and PpRAD51-2 (10 µM). No binding is observed with replicate presented in Fig. S11.

C. Interaction between PpBRCA2-BRC2, -BRC4 and PpRAD51-2. The left panel shows the curves of PpBRCA2-BRC2 and –BRC4 binding at increasing PpRAD51-2 concentrations (0.625 to 10 µM). The right panel presents the Kd calculated by fitting the plateau shift of the association curve as a function of the PpRAD51-2 concentration (steady-state mode).

D. Interaction between PpBRCA2-BRC3G and PpRAD51-2 using increasing PpRAD51-2 concentrations (1 to 20 µM). The Kd is obtained by fitting the plateau shift of the association curve as a function of the PpRAD51-2 concentration (steady-state mode).

Figure S11

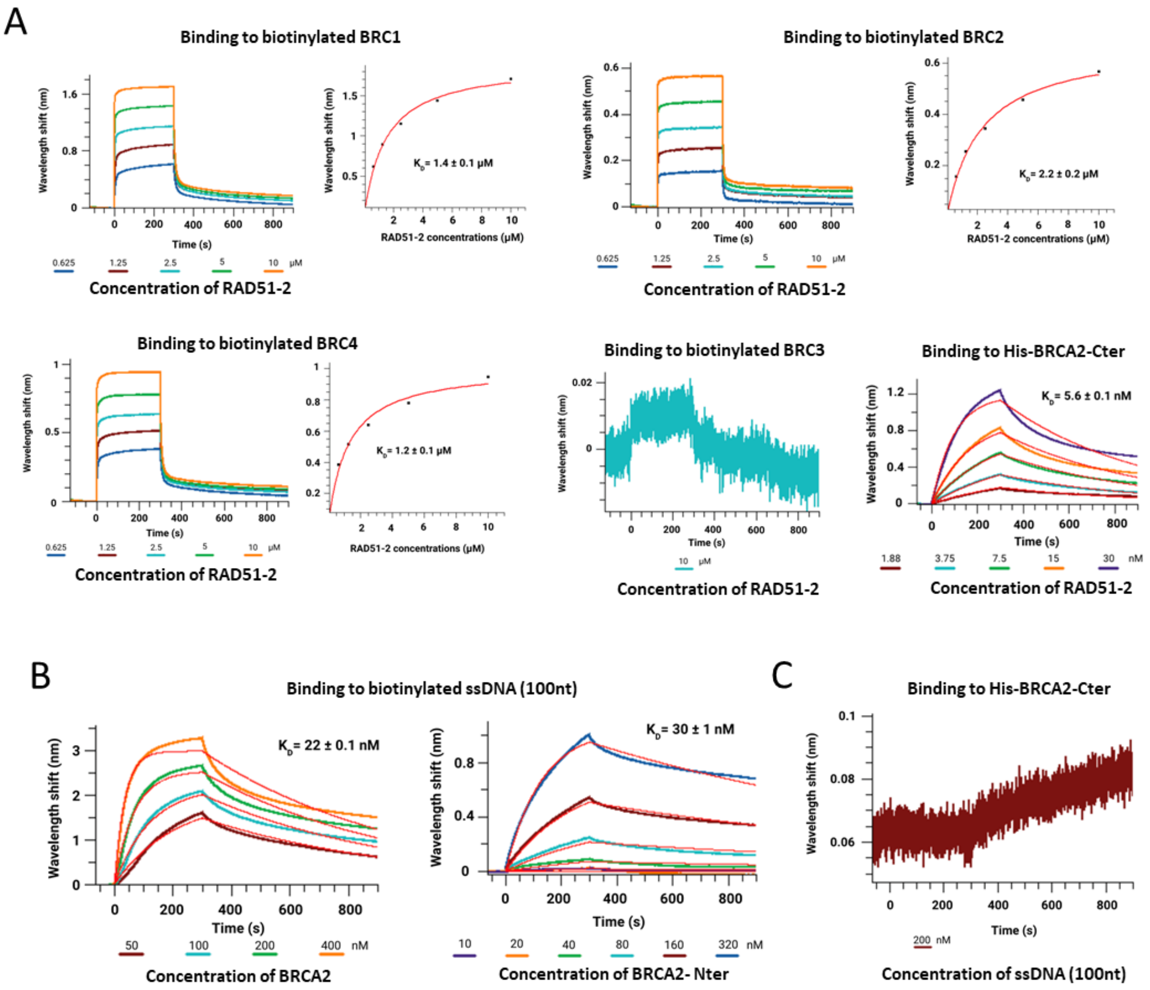


**Figure S11. BRC repeats located in BRCA2-Cter bind to RAD51-2 and BRCA2-Nter binds to ssDNA**

A. Replicates of BioLayer Interferometry (BLI) curves corresponding to the interaction between biotinylated PpBRCA2-BRC peptides and PpRAD51-2.

B. Replicates of BLI curves corresponding to the interaction between biotinylated ssDNA (100 nt) and BRCA2, either full-length, BRCA2-Nter, or BRCA2-Cter.

C. BLI curve corresponding to the interaction between His-PpBRCA2-Cter and ssDNA (100nt).

**DATASET_S1**. HHpred search analysis using Arabidopsis AtBRCA2A as a query on human, drosophila, and moss proteomes.

**DATASET_S2**. HHpred search analysis using PpBRCA2 (Pp6C10_10830) as a query.

**DATASET_S3**. List of all primers, SgRNA, vectors, and peptides, along with raw data from genotoxic and mutator assays.
